# Therapeutic reversal of prenatal pontine ID1 signaling in DIPG

**DOI:** 10.1101/2021.05.10.443452

**Authors:** Viveka Nand Yadav, Micah K. Harris, Dana Messinger, Chase Thomas, Jessica R. Cummings, Tao Yang, Rinette Woo, Robert Siddaway, Martin Burkert, Stefanie Stallard, Tingting Qin, Brendan Mullan, Ruby Siada, Ramya Ravindran, Michael Niculcea, Kevin F. Ginn, Melissa A.H. Gener, Kathleen Dorris, Nicholas A. Vitanza, Susanne V. Schmidt, Jasper Spitzer, Jiang Li, Mariella G. Filbin, Xuhong Cao, Maria G. Castro, Pedro R. Lowenstein, Rajen Mody, Arul Chinnaiyan, Pierre-Yves Desprez, Sean McAllister, Cynthia Hawkins, Sebastian M. Waszak, Sriram Venneti, Carl Koschmann

**Affiliations:** Department of Pediatrics, Division of Pediatric Hematology-Oncology, University of Michigan Medical School; Ann Arbor, USA; Department of Neurology, University of Michigan Medical School; Ann Arbor, USA; Cancer Research, California Pacific Medical Center Research Institute; San Francisco, USA; Arthur and Sonia Labatt Brain Tumour Research Centre, Hospital for Sick Children, University of Toronto; Toronto, Canada; Centre for Molecular Medicine Norway (NCMM), Nordic EMBL Partnership, University of Oslo and Oslo University Hospital; Oslo, Norway; Department of Pediatric Research, Division of Pediatric and Adolescent Medicine, Rikshospitalet, Oslo University Hospital; Oslo, Norway; Department of Computational Medicine and Bioinformatics, University of Michigan Medical School; Ann Arbor, USA; Department of Pediatrics, Children’s Mercy Kansas City; Kansas City, USA; Department of Pathology and Laboratory Medicine, Children’s Mercy Kansas City; Kansas City, USA; Department of Pediatrics, University of Colorado School of Medicine; Aurora, USA; Department of Pediatrics, Seattle Children’s; Seattle, USA; Institute of Innate Immunity, AG Immunogenomics, University Bonn; Bonn, Germany; Department of Pediatric Oncology, Dana-Farber Boston Children’s Cancer and Blood Disorders Center; Boston, USA; Department of Pathology, University of Michigan Medical School; Ann Arbor, USA; Departments of Neurosurgery and Cell and Developmental Biology, University of Michigan Medical School; Ann Arbor, USA

## Abstract

Diffuse intrinsic pontine glioma (DIPG) is a highly aggressive brain tumor with rare survival beyond two years. This poor prognosis is largely due to the tumor’s highly infiltrative and invasive nature. Previous reports demonstrate upregulation of the transcription factor ID1 with H3K27M and *ACVR1* mutations, but this has not been confirmed in human tumors or therapeutically targeted. We developed an in utero electroporation (IUE) murine H3K27M-driven tumor model, which demonstrates increased ID1 expression in H3K27M- and *ACVR1*-mutated tumor cells. In human tumors, elevated ID1 expression is associated with H3K27M/*ACVR1*-mutation, brainstem location, and reduced survival. The *ID1* promoter demonstrates a similar active epigenetic state in H3K27M tumor cells and murine prenatal hindbrain cells. In the developing human brain, ID1 is expressed highest in oligo/astrocyte-precursor cells (OAPCs). These ID1^+^/SPARCL1^+^ cells share a transcriptional program with astrocyte-like (AC-like) DIPG cells, and demonstrate upregulation of gene sets involved with regulation of cell migration. Both genetic and pharmacologic [cannabidiol (CBD)] suppression of ID1 results in decreased DIPG cell invasion/migration in vitro and invasion/tumor growth in multiple in vivo models. CBD reduces proliferation through reactive oxygen species (ROS) production at low micromolar concentrations, which we found to be achievable in the murine brainstem. Further, pediatric high-grade glioma patients treated off-trial with CBD (n=15) demonstrate tumor ID1 reduction and improved overall survival compared to historical controls. Our study identifies that *ID1* is upregulated in DIPG through reactivation of a developmental OAPC transcriptional state, and ID1-driven invasiveness of DIPG is therapeutically targetable with CBD.

**One Sentence Summary:** The transcription factor ID1 is upregulated in a subset of DIPG tumor cells, and ID1-driven invasiveness is therapeutically targetable with CBD.

## INTRODUCTION

Diffuse intrinsic pontine glioma (DIPG) is a lethal pediatric brain tumor that originates in the pons (*1*). With a median survival of 10-11 months, DIPG remains the most aggressive primary brain tumor in children (*2*). Standard of care consists of palliative radiation, and experimental chemotherapies have yet to demonstrate benefit beyond radiation (*2*). Even with the advent of precision-based medicine, clinical trials targeting specific molecular targets are lacking, highlighting the need to identify novel therapeutic targets in DIPG.

As many as 80% of DIPGs harbor mutations in histone H3, which leads to a lysine-to-methionine substitution (H3K27M) in *H3.3A* (*H3F3A*) and *H3C2* (*HIST1H3B*) (*1, 3*). H3K27M is now understood to define a distinct clinical and biological subgroup in DIPG, and is associated with a worse prognosis (*4*). The H3K27M mutation represses the polycomb repressive complex 2 (PRC2), resulting in global reduction of H3K27me3 (with focal gains) (*5*) and global increases in acetylation of H3K27 (H3K27ac), associated with upregulation of tumor-driving genes (*6, 7*).

Basic helix-loop-helix (bHLH) transcription factors are key regulators of tissue and lineage-specific gene expression, and constitutive expression of Inhibitor of DNA binding (ID) proteins have been shown to inhibit the differentiation of multiple tissues (*8*). ID proteins dimerize with bHLH transcription factors, preventing DNA binding (*9*). Overexpression of the Inhibitor of DNA binding 1 (*ID1*) gene has been tied to the pathogenesis of multiple human cancers (*10–12*). A role for ID1 in DIPG has been proposed, based on its downstream association with activin A receptor type 1 (ACVR1) signaling, which is recurrently mutated/activated in 25% of human DIPGs (*13–15*). Germline *ACVR1* mutations in the congenital malformation syndrome fibrodysplasia ossificans progressiva (FOP) activate the bone morphogenetic protein (BMP) signaling pathway, through enhanced recruitment and phosphorylation of SMAD1/5/8, which in turn increases ID1 expression (*16*). Prior studies have shown K27M and *ACVR1* to upregulate *ID1* in cultured human astrocytes and murine models of DIPG (*13, 14*). ID1 has been shown to drive an invasive tumor phenotype in multiple solid tumors (*10, 11*). Invasion into normal pontine tissue is a pathognomonic feature of DIPG, but its regulation remains poorly understood. Further, analysis of ID1 in human DIPG, and its regulation and targetability, have not been previously investigated.

In the present study, we show that human DIPGs demonstrate epigenetic activation and increased expression of ID1, influenced by H3K27M and *ACVR1* mutational status and brain location. This epigenetic activation mimics ID1 regulation in the developing human and murine prenatal pons. Genetic knockdown and pharmacologic [cannabidiol (CBD)] inhibition of ID1 decreases invasion and migration and improves survival in multiple preclinical DIPG models and human patients. These findings represent an exciting new direction for understanding the regulation and targetability of invasion in DIPG, with broad implications for therapeutic targeting of solid tumors with ID1 up-regulation.

## RESULTS

### Increased ID1 expression with H3K27M and ACVR1 mutations in murine DIPG tumor model

We first sought to confirm whether ID1 expression is affected by the presence of H3K27M and *ACVR1* mutations. We adopted an in utero electroporation (IUE) model of pediatric high-grade glioma (pHGG), as previously described by our group (*17*). Mice developed tumors [mutant T<u>P</u>53, mutant <u>P</u>DGFRA (D842V) with *H3.3A* <u>K</u>27M mutation (“PPK”) or *H3.3A* <u>w</u>ildtype (“PPW”)] via plasmid injection into the lateral ventricles of E13.5 embryonic CD1 mice (Fig. 1A-B). Transfection efficiency and tumor growth/size were monitored using in vivo bioluminescence imaging, and primary neurosphere cell cultures were generated for each group by tumor dissociation (Fig. 1B). Survival analysis revealed that PPK mice (n=15) had significantly reduced survival compared to their H3^Wildtype (WT)^ counterparts (PPW; n=10) (Fig. 1C). Additionally, immunohistochemistry (IHC) and western blot analyses of murine tumors showed tumor-specific expression of H3K27M and global loss of H3K27me3 expression, a salient feature expected in H3K27M-mutant DIPG tumors (Fig. 1D-E) (*18*). Importantly, ID1 expression was elevated in PPK tumors compared to PPW (Fig. 1E). In order to determine the impact of *ACVR1* mutation on ID1 expression in DIPG, we introduced *ACVR1* mutation via lentiviral (LV) transduction into PPK tumor cells and primary *H3.3A* K27M/*ACVR1*^WT^ human DIPG cells (DIPGXIIIp). Western blot analysis revealed increased ID1 expression and SMAD activation with the introduction of *ACVR1* mutation in both PPK and DIPGXIIIp tumor cells (Fig. 1F), consistent with previous reports (*13, 14*).

**Fig. 1.**
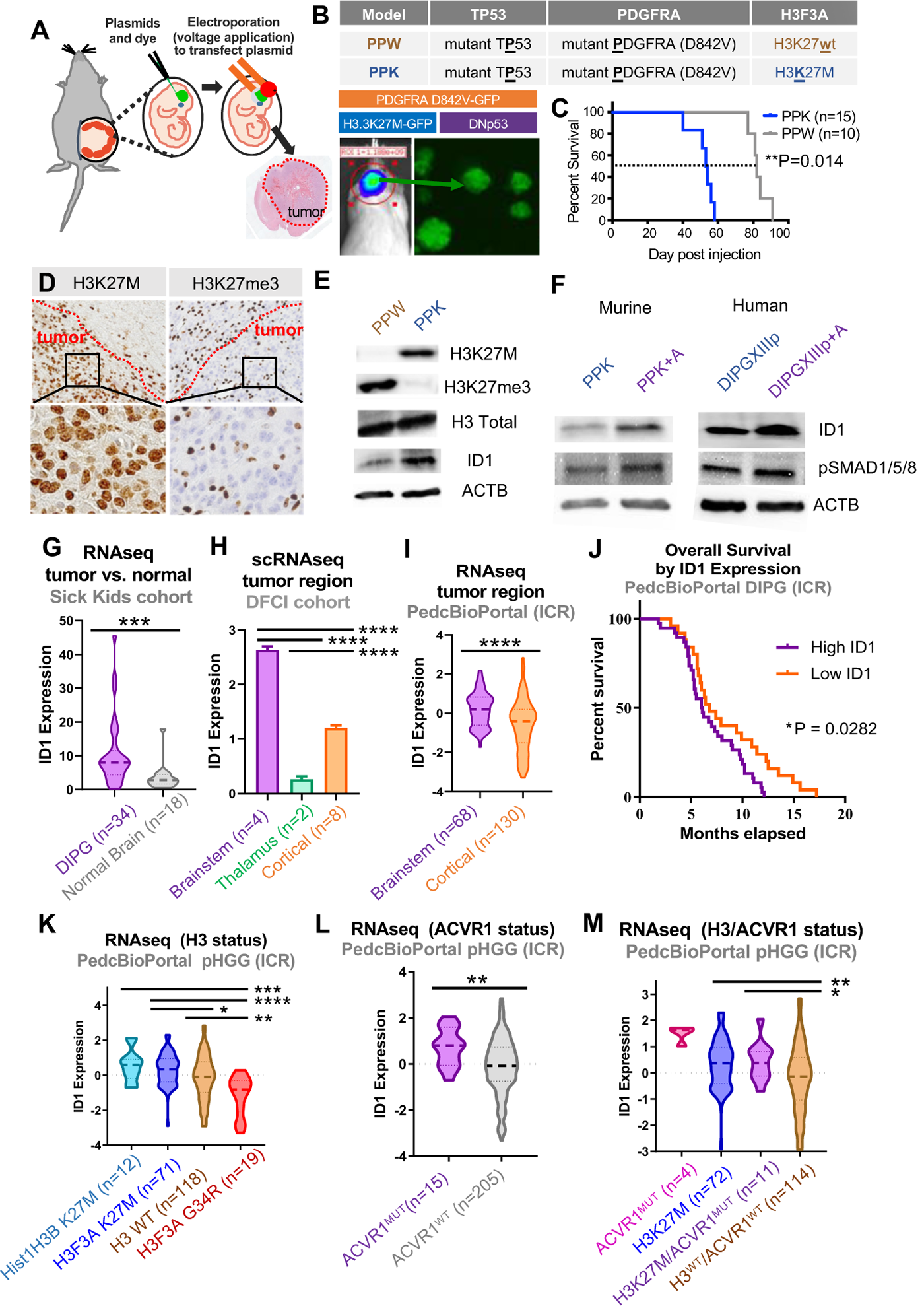
Elevated expression of *ID1* in DIPG population. **(A)** IUE-mediated H3K27M-tumor model. **(B)** PDGFRA^MUT^-p53^MUT^-H3^WT^ (“PPW”) and PDGFRA^MUT^-p53^MUT^-H3K27M (“PPK”) tumors are generated by IUE. Tumor growth is monitored by in vivo bioluminescence imaging and primary neurosphere cell cultures are generated by dissociation of tumor tissue. **(C)** Survival curve for PPW and PPK mice displays significantly reduced survival of PPK compared to PPW mice; **P=0.014, log-rank test. **(D)** IHC-stained images of PPK tumor show tumor-specific H3K27M expression and reduced H3K27me3 (representative of n=3 PPK tumors). Magnification = 10x (top row); 40x (bottom row). **(E)** Western blot (WB) of PPW and PPK primary neurospheres for assessment of H3K27M, H3K27ac and ID1 expression by H3 mutational status. **(F)** WB of murine PPK and PPK+*ACVR1*^MUT^ (“PPK+A”) cells, and human DIPGXIIIp and DIPGXIIIp+*ACVR1*^MUT^ cells, for assessment of ID1 and pSMAD expression. **(G)** *ID1* expression of DIPG tissue (n=34) compared to matched normal brain tissue (n=18) from the SickKids cohort; ***P < .001, unpaired parametric t-test. **(H)** *ID1* expression by scRNA-seq from the DFCI cohort, including brainstem (n=4), thalamus (n=2) and cortex (n=8); ****P<0.0001, one-way ANOVA t-test. **(I)** *ID1* expression of DIPG tissue (n=68) compared to hemispheric pHGG tissue (n=130). Data from ICR cohort; ****P < 0.0001, unpaired t-test. **(J)** Kaplan-Meier survival curve of DIPG patients (n=66) grouped by high and low *ID1* expression. *P =0.0282, Mantel-Cox test. **(K)** *ID1* expression across *Hist1H3B (H3C2)* K27M (n=12), *H3F3A (H3.3A)* K27M (n=71), H3^WT^ (n=118) and *H3F3A (H3.3A)* G34R (n=19) DIPG tumors. Data from ICR cohort, presented in Mackay et al; *P<0.05, **P<0.01, ***P<0.001, ****P<0.0001, one-way ANOVA t-test. **(L)** *ID1* expression of pHGG tissue by *ACVR1* mutational status (n=15 ACVR1^MUT^; n=205 ACVR1^WT^). Data from ICR cohort; **P<0.01, unpaired parametric t-test. **(M)** *ID1* expression of pHGG tumors with *ACVR1* mutation only (n=4), H3K27M only (n=72), H3K27M and *ACVR1* mutations (n=11) and neither mutation (H3WT/*ACVR1* WT; n=114). Data from ICR cohort; *P<0.05, **P<0.01, one-way ANOVA t-test.

### ID1 expression increased in human DIPG and associated with lower overall survival

We next assessed the impact of H3K27M (*H3.3A* or *H3C2*) and *ACVR1* mutations on *ID1* expression in DIPG and non-brainstem pHGG. Whole transcriptome sequencing was performed on 34 DIPG and 18 normal post-mortem brain tissue specimens taken from a single institutional cohort (Sick Kids, Toronto). Compared to normal brain (cortex), DIPG tissue showed significantly higher *ID1* expression (Fig. 1G). Single cell RNA-sequencing (scRNA-seq) data from H3K27M-mutant DIPG tumors [Dana-Farber Cancer Institute (DFCI) cohort (*19*)] confirmed that malignant cells display significantly higher *ID1* expression compared to nonmalignant cells within these tumors (Fig. S1A). ScRNA-seq data from H3K27M-pHGG patients (n=14) revealed higher *ID1* expression in pontine H3K27M-DIPG cells compared to thalamic and cortical pHGG tumors (Fig. 1H). This was confirmed in bulk RNA-seq [ICR cohort (Institute for Cancer Research), n=198 (*20*)], in which brainstem pHGG tumors (DIPG) showed significantly higher *ID1* expression than cortical pHGGs (Fig. 1I). High *ID1* expression has been linked to lower overall survival (OS) in multiple cancers (*21*). Indeed, DIPG patients with higher bulk *ID1* expression (ICR cohort) have lower OS (Fig. 1J). These data support that ID1 is involved in the pathogenesis of human DIPG.

### ID1 expression influenced by H3 and ACVR1 mutational status in human DIPG

Introduction of the recurrent mutations *H3.3A* K27M and *ACVR1* have been shown to increase *ID1* expression in cultured astrocytes (*13, 14*), consistent with findings in our IUE tumor model (Fig. 1E). Analysis of bulk tumor RNA-seq (ICR cohort) revealed that *ID1* expression is significantly increased in pHGGs harboring H3K27M (*H3.3A* or *H3C2*) compared to H3^WT^ and H3G34R tumors (Fig. 1K) (*20*). *ACVR1*-mutant tumors (Fig. 1L) and those with co-mutation (H3K27M and *ACVR1*, Fig. 1M) have significantly higher *ID1* expression compared to WT tumors. Interestingly, in scRNA-seq data (DFCI cohort), elevated *ID1* expression is seen in a higher proportion of malignant cells within pontine H3K27M tumors (n=4; 35-69%) in comparison to thalamic H3K27M tumors (n=2; 6-9%) (Fig. S1B). Taken together, these data support that *ID1* expression in pHGG is driven by both mutational status of H3 and *ACVR1* and regional (anatomic) influences.

### Epigenetic state of ID1 loci in H3K27M tumor cells and murine prenatal hindbrain cells

In patients with germline *ACVR1*-mutant FOP or DIPG tumors with somatic *ACVR1* mutations, ID1 expression is activated by BMP signaling (*15, 22*). However, the mechanism of H3K27M mutation promoting increased ID1 expression is less understood. We assessed whether H3K27ac and H3K27me3 marks at regulatory regions of the *ID1* gene could be contributing to the increased *ID1* expression observed in human DIPG. Quantitative PCR (qPCR) demonstrated *ID1* expression to be higher in H3K27M and *ACVR1*-mutant DIPG autopsy samples (n=4 tumor sites) compared to H3WT/*ACVR1*^WT^ DIPG tissue (n=6 tumor sites) and normal brain tissue samples (n=4 sites) Fig. 2B and S2A-C). ChIP-Seq at the ID1 gene loci on normal adolescent pontine (n=1), H3^WT^ DIPG (n=1) and H3K27M DIPG (n=4) samples revealed a marked increase in H3K27ac deposition at *ID1* gene body elements in H3K27M DIPG tumor tissue compared to H3^WT^ DIPG tumor and normal pontine tissue, with minimal H3K27me3 marks across the ID1 loci in all tissue types (Fig. 2C). Subsequent ChIP-qPCR for quantification (primers in Supplemental Table 1) demonstrated significantly elevated H3K27ac at predicted promotor and gene body regions of the ID1 locus compared to H3WT/ACVR1^WT^ DIPG tumor samples (Fig. 2D). Decreased H3K27me3 was also observed, though this was only significant at one of the predicted promotor regions between H3K27M/ACVR1^MUT^ and H3^WT^/ACVR1^WT^ DIPG sample groups (Fig. 2E). Taken together, however, the effects of these changes in H3K27ac and K3K27me3 marks correspond with H3K27M-mutant samples being epigenetically activated for ID1 expression.

**Fig. 2.**
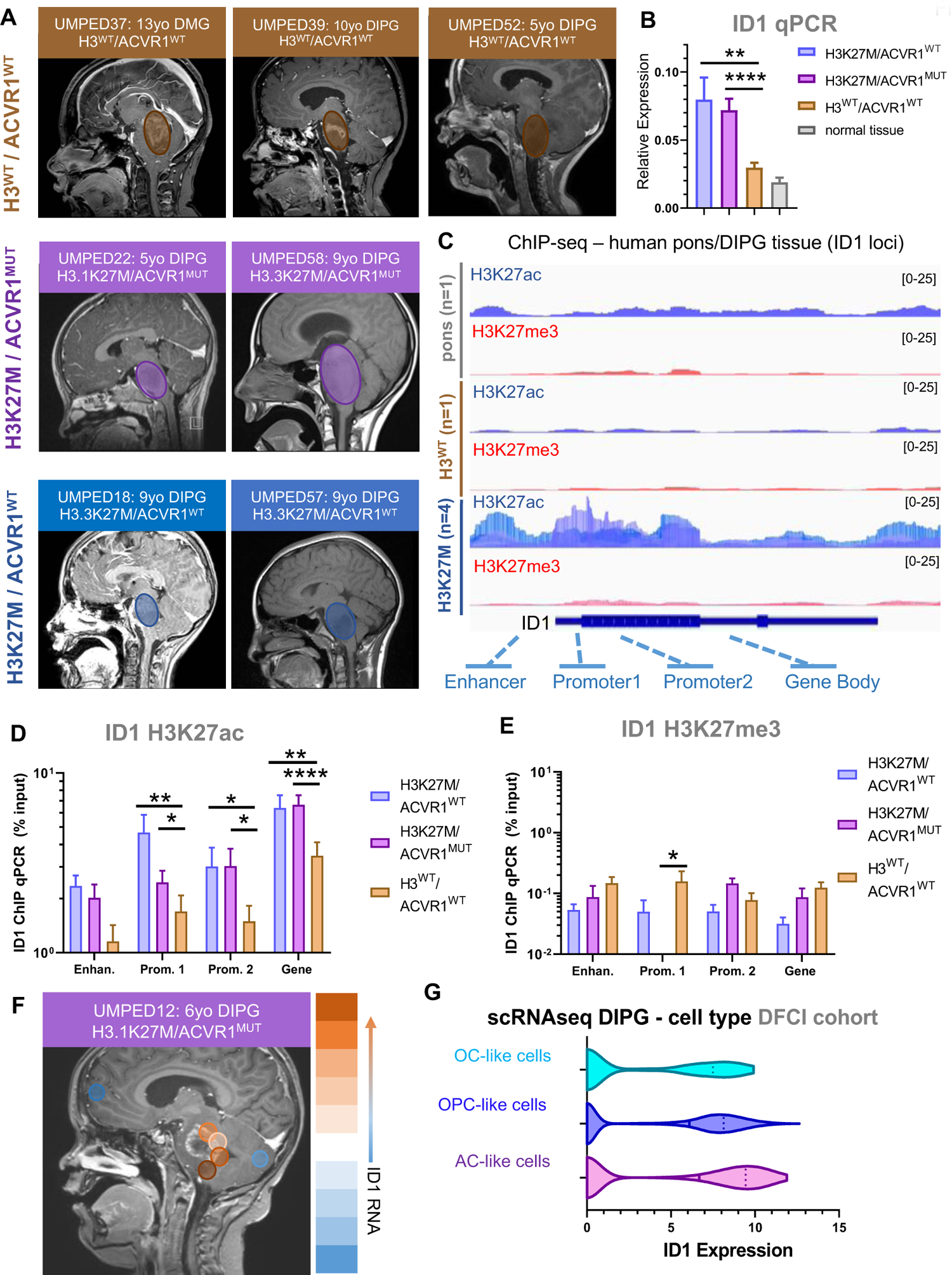
*ID1* is epigenetically active in H3K27M-DIPG. **(A)** Multifocal DIPG tumor samples were obtained at autopsy from n=2 patients with H3K27M mutation and wildtype *ACVR1* (ACVR1^WT^), n=2 patients with H3K27M mutation and *ACVR1* mutation (ACVR1^MUT^) and n=3 patients with wildtype H3 and *ACVR1*. Circles over MRI images represent the approximate region of tumor. **(B)** *ID1* expression (qPCR) for multifocal samples collected from patients in (A). Data represent mean+/-SEM; **P<0.01, ****P<0.0001, one-way ANOVA t-test. **(C)** ChIP-sequencing of H3K27ac and H3K27me3 deposition at the *ID1* gene locus in normal human pontine tissue (n=1), H3^WT^ DIPG tumor tissue (n=1) and H3K27M DIPG tumor tissue (n=1). **(D-E)** ChIP-qPCR quantification of deposited (D) H3K27ac, and (E) H3K27me3 marks at gene body elements identified in part C for the *ID1* gene. Data represent samples from patients in (A), mean+/-SEM; *P<0.05, **P<0.01, ****P<0.0001, one-way ANOVA t-test. **(F)** MRI image of H3K27M/ACVR1^MUT^ DIPG patient with circles representing regions where samples were obtained at autopsy. Color scale on right displays relative level of *ID1* expression by qPCR (orange=higher *ID1* expression; blue=lower *ID1* expression. **(G)** ScRNA-seq data (DFCI, n=4 DIPGs) of malignant DIPG cells plotted to show ID1 expression across varying subtypes of cells [oligodendrocyte-like (OC-like); OPC-like; AC-like].

While brainstem tumors broadly show increased *ID1* expression compared to normal brain, we noted differences in expression by qPCR between multi-focal autopsy samples. Expanded multi-focal (n=6) bulk RNA-sequencing of a single H3K27M/*ACVR1*-mutant DIPG patient (UMPED12) confirmed varying levels of *ID1* expression across different regions of the tumor (Fig. 2F). This finding led us to analyze scRNA-seq in order to determine whether a specific malignant cell subpopulation could be contributing to the increased *ID1* expression seen in DIPG. Assessment of *ID1* expression across all malignant cell types in DIPG cells from four patients showed that *ID1* is most highly expressed in DIPG cells with an astrocytic differentiation program [“AC-like cells” (*19*)], followed by oligodendrocyte precursor cell-like (“OPC-like”) cells (Fig. 2G and S3A). OPC-like cells are known to constitute the majority of cycling cells in DIPG (*19*). Previous analysis showed that nearly all cycling DIPG cells have an OPC-like phenotype (*19*) and we observed higher levels of *ID1* expression in cycling compared to non-cycling cells (Fig. S3B).

### Single-cell transcriptional analysis of ID1^+^ cells in human developing brain and H3K27M tumors

Anatomic location and developmental context strongly influence the formation of many pediatric tumors, including DIPG. We next assessed *ID1* expression and histone modifications across pre- and post-natal mouse brain developmental stages. RNA in-situ hybridization data (Allen Brain Atlas) demonstrated *ID1* to be highest expressed in the developing prenatal mouse hindbrain (including the developing pons) compared to forebrain or midbrain, with minimal *ID1* expression throughout the entire postnatal mouse brain (Fig. 3A-B and S4). In E15.5 mouse brains, ENCODE data (*23, 24*) revealed H3K27ac to be elevated at *ID1* enhancer sites in the hindbrain compared to midbrain and forebrain (Fig. S5A-B).

**Fig. 3.**
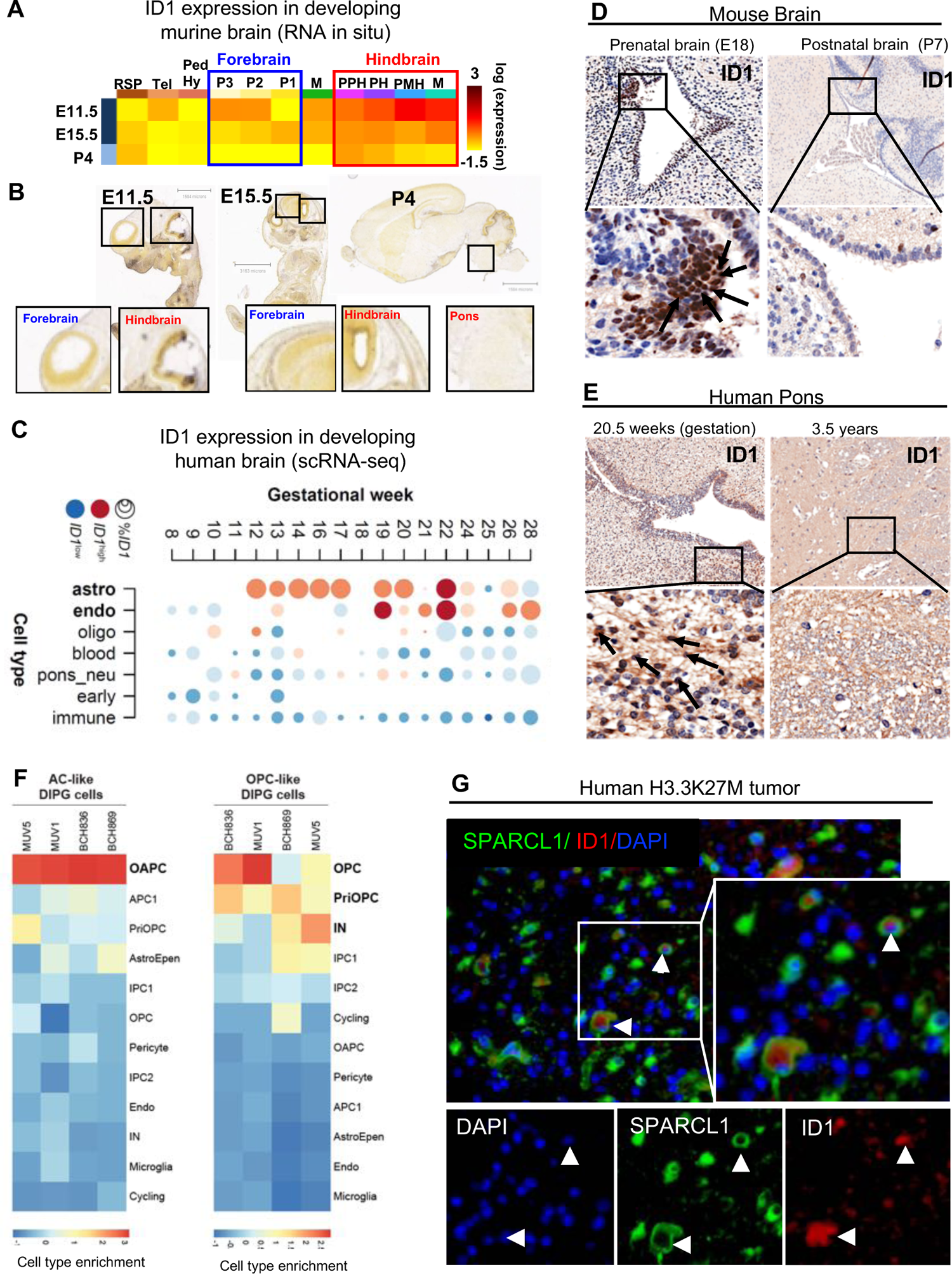
ID1 expression is elevated in developing astrocyte cells in prenatal human and murine hindbrain. **(A)** Heat map showing relative *ID1* expression by in situ hybridization (ISH) in murine brain across development. Available from: http://developingmouse.brain-map.org/. **(B)** ISH of sagittal developing murine brain sections showing higher *ID1* RNA in embryonic hindbrain than forebrain, and minimal *ID1* RNA in all post-natal brain [Allen Developing Mouse Brain Atlas. Available from: http://developingmouse.brain-map.org/]. **(C)** Heatmap of *ID1* expression across varying cell types during normal human pontine development [data from Fan et al. (*25*)]. Circle size indicates the percentage of cells that express ID1 and color indicates the expression level in ID1^+^ cells (red=high expression; blue=low expression). **(D)** ID1 IHC staining of normal human pontine tissue displays higher ID1 expression in cells lining the 4^th^ ventricle at 20.5 weeks gestation and minimal expression in brain tissue at 3.5 years of age. **(E)** ID1 IHC of normal murine pontine tissue at embryonic day 18 (E18) displays higher ID1 expression compared to postnatal day 7 (P7). Magnification = 10x (top row); 40x (bottom row). **(F)** Overlap of genes expressed by cell types in the developing human pons Fu et al. (*28*) in DIPG tumor cell subsets. (Red=cell type marker genes enriched in DIPG cells; blue=cell type marker genes not enriched in DIPG cells). **(G)** Immunostaining of SPARCL1 (green) and ID1 (red) in human DIPG tissue showing co-localization of ID1 and SPARCL1 in a subset of cells (white arrow). Scale bar, 20 µm. Tumor nuclei were stained with DAPI (blue). [For (A), from left to right (row headings), RSP: rostral secondary prosencephalon, Tel: telencephalic vesicle, PedHy: peduncular (caudal) hypothalamus, P3: prosomere 1, P2: prosomere 2, P1: prosomere 3, M: midbrain, PPH: prepontine hindbrain, PH: pontine hindbrain, PMH: pontomedullary hindbrain, MH: medullary hindbrain (medulla); from top to bottom (column headings), E11.5/15.5: embryonic day 11.5/15.5, P4: postnatal day 4].

Analysis of developing human (*25*) and mouse (*26*) brain scRNA-seq data showed that *ID1* expression peaks at gestational week (GW) 12-22 in the human pons (Fig. 3C) and early postnatal mouse pons (P0; Fig. S6-S7), and is most highly expressed in astrocytes. *ID1* expression is also high in human endothelial cells, consistent with previous data (Fig. 3C) (*27*). IHC analyses of pre- and post-natal brains confirmed elevated ID1 in the murine embryonic brain (E18; Fig. 3D) and human GW 20.5 brain (Fig. 3E) in subventricular regions lining the 4^th^ ventricle, compared to all postnatal brain locations.

We next sought to assess whether *ID1^+^* sub-populations of malignant DIPG cells share a transcriptional program with *ID1^+^* developing brain cells. Interestingly, AC-like cells from all four DIPG tumors show the strongest overlap with the transcriptional program of the recently defined OAPC cell population (*28*) in the developing human brain (Fig. 3F). The OAPC program was not enriched in OPC-like cells in any of the four DIPG tumors (Fig. 3F). OAPCs are present primarily in the outer subventricular zone during the neurogenesis-to-gliogenesis switch period and express both astrocyte (GFAP) and oligodendrocyte (OLIG1, OLIG2) marker genes as well as SPARCL1, which is involved in regulation of cell adhesion (*28*). We found *ID1* to be a marker gene for both AC-like DIPG cells and OAPCs. Immunofluorescence of human H3K27M-DIPG samples revealed co-localization of ID1 and SPARCL1 expression in sub-populations of cells (Fig. 3G). Assessment of SPARCL1 expression across all malignant cell types in DIPG cells from four patients showed that SPARCL1 is most highly expressed in AC-like DIPG cells. Importantly, AC-like DIPG cells demonstrate enrichment of gene sets involved in regulation of cell adhesion and migration (Fig. S9), further implicating the potential role of *ID1*^+^ AC-like cells in the regulation of DIPG tumor cell invasion and migration.

### Impact of genetic and pharmacologic knockdown of ID1 on invasion and migration

To examine the phenotypic impact of *ID1* in human DIPG cells, a patient-derived DIPG cell culture with *H3.3A* K27M and *ACVR1* mutation (DIPG007) was lentivirally-transduced with ID1-targeting shRNA or scrambled shRNA control (Fig. 4A). *ID1* knockdown (shRNA-64) resulted in reduced SPARCL1 expression in DIPG007 cells by western blot, further implicating the role of *ID1* in the regulation of this OAPC/AC-like cell marker gene (Fig. 4B). *ID1* knockdown significantly reduced DIPG007 invasion (Fig. 4C) and migration, as measured by scratch assay percent wound closure (Fig. 4D-E). In comparison, invasion and migration of human embryonic kidney cell line HEK293 was not affected upon *ID1* knockdown (Fig. S10A-C).

**Fig. 4.**
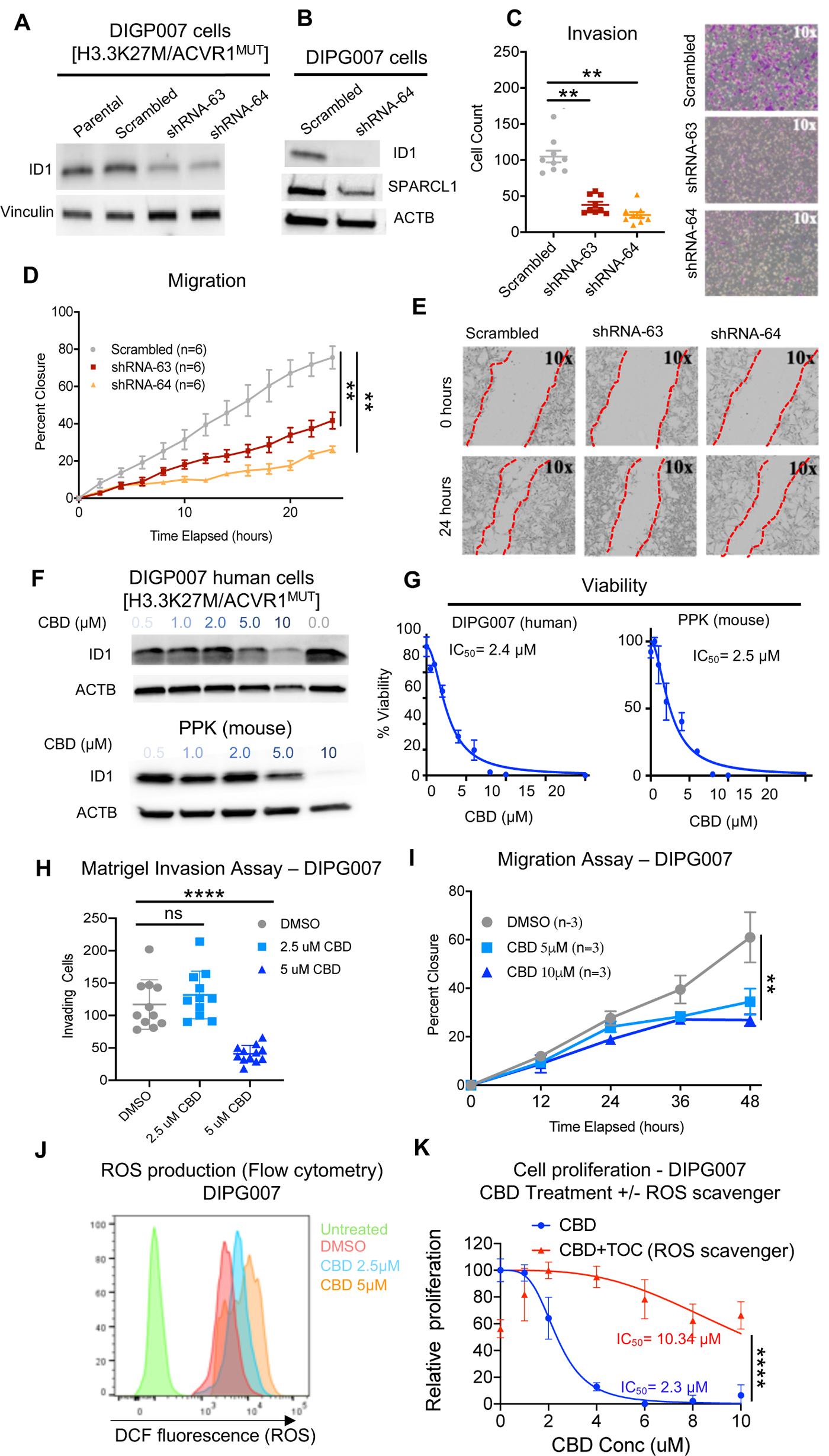
Genetic and Pharmacological inhibition of *ID1* decreases DIPG cell invasion and migration. **(A)** Western blot (WB) confirming *ID1* knockdown in DIPG007 cells. (B) WB depicting reduction in SPARCL1 expression along with decreased ID1 expression in *ID1*-knockdown DIPG007 cells. **(C)** Effect of *ID1* knockdown on invasion as measured by Matrigel-coated Boyden chamber assay. Images show invading cells stained with crystal violet. Each data point represents an individual image; **P < 0.01, unpaired parametric t-test. **(D-E)** Effect of *ID1* knockdown on DIPG007 migration as measured by scratch assay, quantified as percent wound closure. Images show representative scratch at 0 and 24 hours outlined in dotted red line. Experiment was completed in triplicate and data points represent mean+/-SEM, **P < 0.01; images taken with Incucyte; area measured by ImageJ. **(F)** WB for ID1 and ACTB expression in DIPG007 and PPK cells treated with increasing concentrations of CBD or DMSO control. **(G)** Viability of DIPG007 and PPK cells treated with increasing concentrations of CBD (0.5-20µM) relative to DMSO-treated control. Experiment was completed in triplicate and data points represent mean+/-SEM. **(H)** DIPG007 cells were treated for 2 days with DMSO (control), 2.5µM or 5µM CBD and invasion was measured by Matrigel-coated Boyden chamber. Each data point represents an individual image, mean+/-SEM; ****P < 0.0001, unpaired parametric t-test. **(I)** Effect of CBD treatment (5-10µM) on DIPG007 migration as measured by scratch assay, quantified as percent wound closure. Experiment was completed in triplicate and data points represent mean+/-SEM, **P < 0.005, two-way ANOVA t-test. **(J)** Histogram showing increase in DCF (ROS) with increasing doses of CBD. **(K)** Production of ROS mediates the inhibitory activity of CBD through ID1. DIPG007 cells were treated for 72 hours with vehicle (DMSO) or different concentrations of CBD (10, 8, 6, 4, 2, 1 µM) in the presence and absence of 50 µM TOC. IC_50_ was 2.3µM for CBD treatment alone and 10.34µM for CBD + TOC; ****P < 0.0001, two-way ANOVA t-test. Cell proliferation was measured using XTT assay.

A few compounds that reduce *ID1* expression include Cannabidiol (CBD), Pimozide, 2-Methoxyestradiol and MK615 (*29–31*). Of these, CBD is the most studied, clinically available and CNS-penetrant agent (*32, 33*). CBD is the non-psychoactive compound found in *Cannabis sativa* (*34*). CBD has wide-ranging impacts on cellular behavior, including the ability to downregulate expression of *ID1* and to inhibit invasion in multiple pre-clinical cancer models (*12, 35–37*). Based on these studies, we sought to investigate the targeting of ID1 in DIPG through use of CBD. Treatment of human DIPG007 and mouse PPK cells with CBD reduced *ID1* expression (Fig. 4F) and cell viability (Fig. 4G), with an IC_50_ of 2.4 and 2.5 µM, respectively. We also treated two additional human DIPG cell cultures with H3K27M/ACVR1^WT^ status, DIPGXIIIp and PBT-29, with CBD, and found reductions in cell viability at an IC_50_ of 6.8 and 7.2 µM, respectively (Fig. S11A-B). Additionally, CBD treatment resulted in significantly reduced invasion and migration of human DIPG007 cells (Fig. 4H-I and S12A-B) and human PBT-29 cells (Fig. S12C-F) in the 5-10 uM range.

CBD has been reported to increase intracellular levels of reactive oxygen species (ROS) (*38*). In line with this, our data reveal that DIPG007 cells treated with CBD show a dose dependent increase in ROS levels (Fig. 4J). Additional treatment with α-tocopherol (TOC), a ROS scavenger (*37, 38*), severely restricted the ability of CBD to inhibit proliferation of DIPG007 cells (Fig. 4K).

### Genetic knockdown of ID1 in IUE murine model

In order to assess whether *ID1* suppression would impede tumor growth in PPK mice, we developed a PBase-responsive ID1-shRNA plasmid and scrambled short hairpin (“sh-control”). PPK-ShID1 mice exhibited significantly prolonged survival when compared to PPK-Sh-control mice (Fig. 5A). PPK-ShID1 mice demonstrated significantly-extended median survival (p=0.01) and reduced luminescent tumor signals when compared to control mice (Fig. 5B-C). IHC analysis of moribund tumors demonstrated reductions in ID1 and Ki67 expression (Fig. 5D-E) in PPK-ShID1 tumors. PPK-ShID1 tumors also exhibited more distinct tumor borders (e.g. reduced tumor invasion into normal brain) in vivo (Fig. 5F). Implantation of DIPG007 cells with ShID1 (or control) into the brainstem of NSG mice also demonstrated reduced pace of luminescent growth (Fig. S13A-C), although this did not affect overall tumor survival. These data indicate that genetic ID1 knockdown inhibits tumor growth in vivo and reduces tumor invasion and proliferation.

**Fig. 5.**
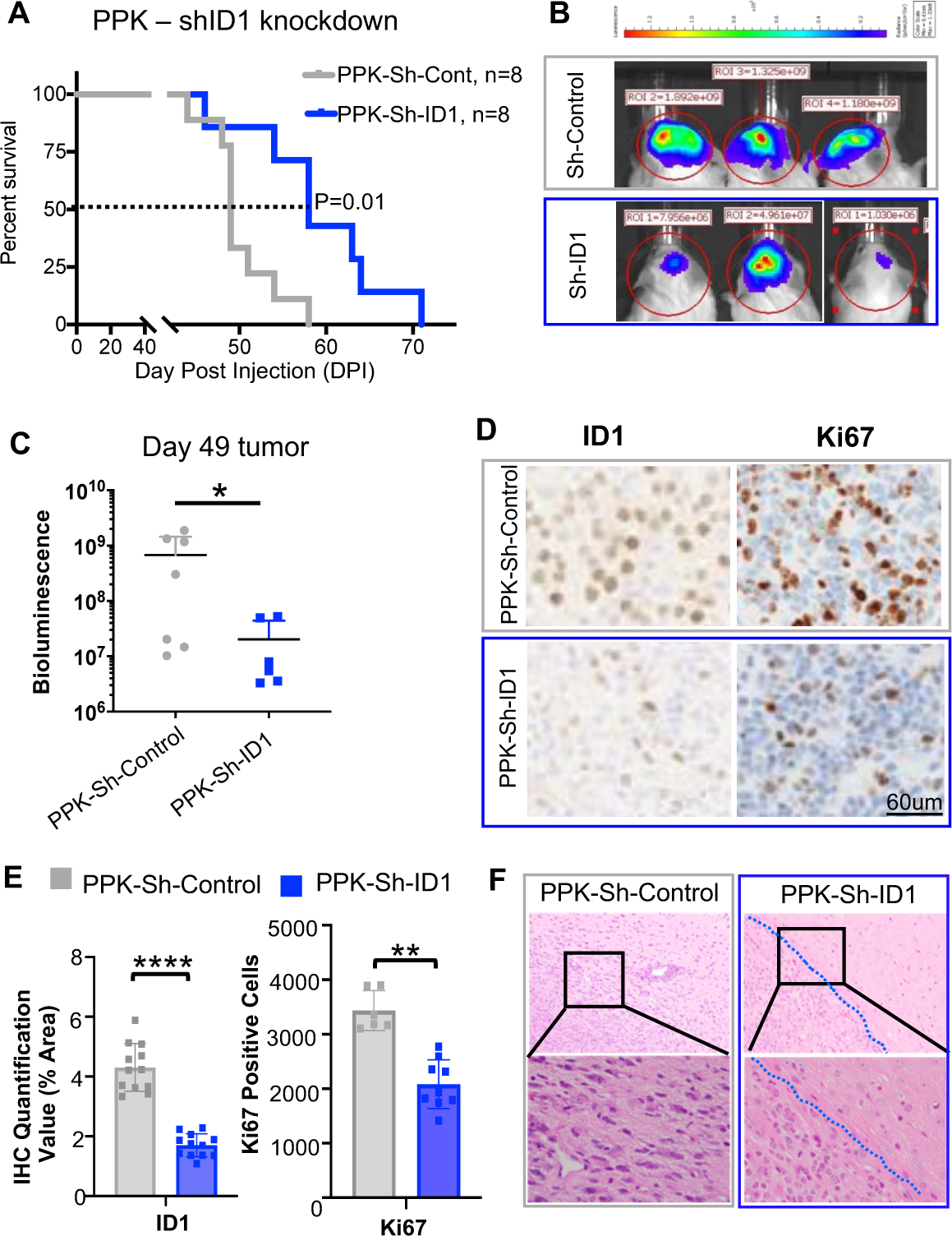
Genetic targeting of ID1 decreases cell viability and slows murine tumor growth in PPK IUE tumor model. **(A)** Standard Kaplan-Meier survival plot reveals notable increase in survival for PPK-Sh-ID1 [*PDGFRA-*, *TP53-* and H3K27M-mutant with ID1 knockout (n=8)] mice with median survival 58 days post-IUE injection compared to PPK-Sh-control (n=8) mice with median survival of 49 days; P=0.01, Log-rank test. **(B)** Representative bioluminescence images of PPK-Sh-control tumors and PPK-Sh-ID1 (representative from n=8), 49 days after IUE injection, displaying lower average luminescence in the PPK-Sh-ID1 group than in the PPK-Sh-control. **(C)** IUE PPK bioluminescence tumor monitor growth data with statistical significance between PPK-Sh-control and PPK-Sh-ID1 groups 49 days after IUE injection. *P<0.05, one-way ANOVA t-test. (**D**) IHC analysis of ID1 and Ki67 expression in tumors from PPK-Sh-ID1 and PPK-Sh-control mice. Images representative of each experimental cohort. Magnification=40x. **(E)** IHC quantification for PPK-Sh-control and PPK-Sh-ID1 mice for ID1 and Ki67 expression levels. **P=0.0065 and ****P ≤ 0.0001, one-way ANOVA t-test. Data points include 3 animals per treatment group and 4 images per animal. Data represent the mean+/-SEM. **(F)** Images of IUE-generated PPK-Sh-Control and PPK-Sh-ID1 tumor borders for assessment of tumor cell invasiveness. Magnification = 10x (top row); 40x (bottom row).

### Pharmacological inhibition of ID1 with Cannabidiol (CBD) in IUE murine model

We next proceeded to our IUE PPK murine model to assess the impact of CBD in vivo. We performed daily treatment with CBD (15 mg/kg), or vehicle control. CBD treatment significantly improved median survival compared to vehicle control (Fig. 6A). Moribund tumors treated with CBD showed reductions in ID1 and Ki67 expression following CBD treatment (Fig. 6B-C). Additionally, CBD-treated tumors displayed reduced invasiveness of tumor cells compared to vehicle-treated mice (Fig. S14). Both genetic (ShID1) and pharmacologic (CBD) knockdown of ID1 in murine models resulted in reduced tumor infiltration into the contralateral hippocampus compared to controls (Fig. 6D). These data indicate that CBD reduces ID1 expression and tumor invasion and significantly improves survival of H3K27M-mutant tumors in vivo.

**Fig. 6.**
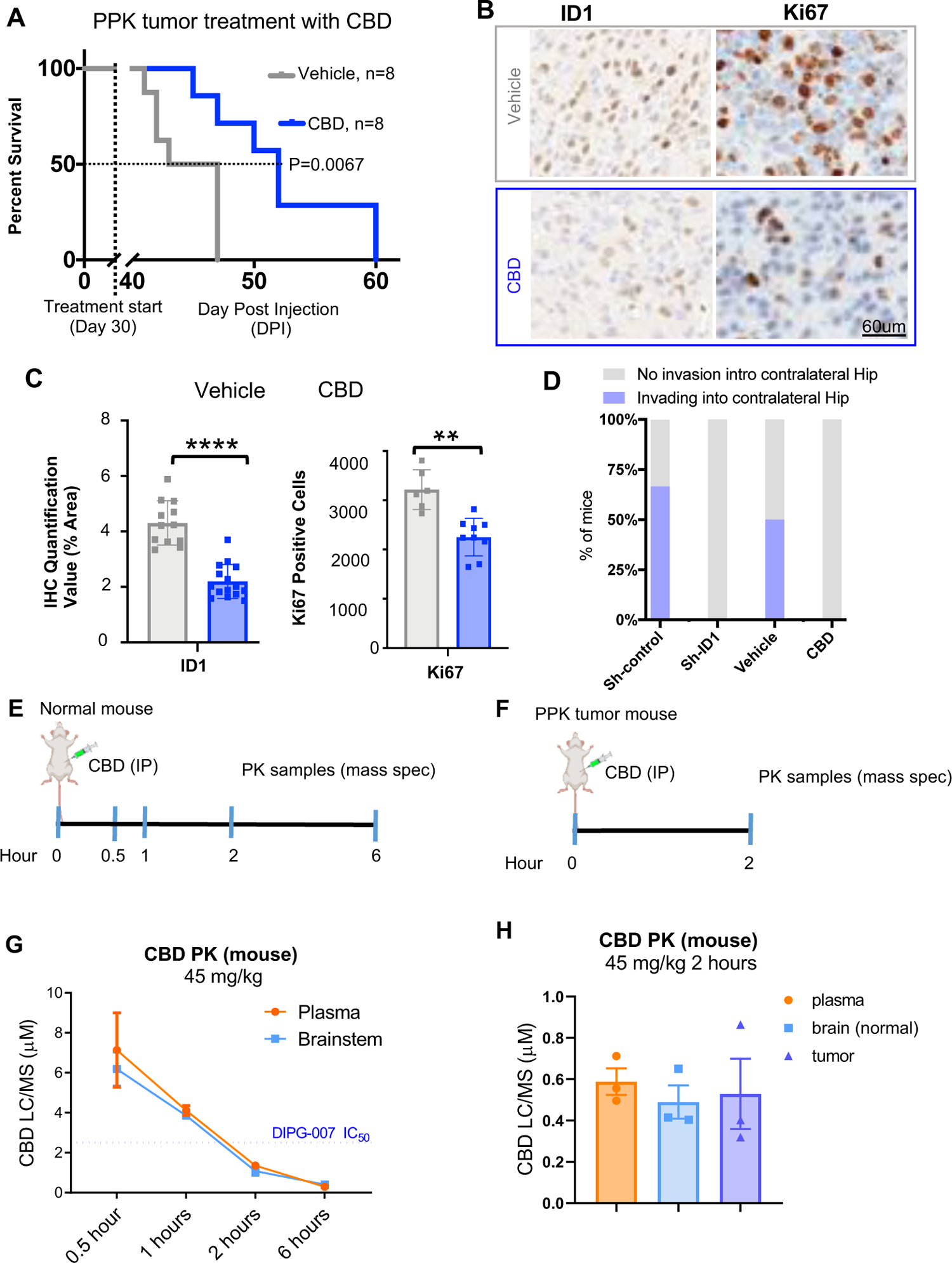
Therapeutic inhibition of ID1 with CBD decreases ID1 expression and improves survival of PPK tumor-bearing mice. **(A)** Survival curve for PPK mice shows that median survival for control condition was 45 days post-IUE injection (n=8) and 55 days for CBD condition (15 mg/kg, n=8). **P<0.005, Log-rank test. **(B-C)** IHC analysis and quantification of tumor images reveals that CBD treatment reduced expression of ID1 and Ki67 compared to vehicle-treated tumors (representative of n=3 tumors); **P=0.0065 and ****P ≤ 0.0001, Dunnett’s multiple comparisons test. N=3 animals per treatment group and 4 images per animal. Data represent the mean+/-SEM. Magnification=10x. **(D)** Analysis of tumor invasion in tumor-bearing mice (n=3 mice per group) with genetic (sh-ID1) or pharmacologic (CBD) ID1 knockdown. Invasion was defined as tumor infiltration into the contralateral hippocampus (Hip). **(E)** Timeline for pharmacokinetic (PK) liquid chromatography (LC)/mass spectrometry (MS) analysis of CBD treatment by intraperitoneal (IP) injection in normal mouse plasma and brainstem. **(F)** Timeline for PK mass spec analysis of CBD treatment by IP injection in PPK mouse plasma, normal brain and tumor. **(G)** PK analysis results for normal (non-tumor-bearing) mice treated with 45 mg/kg CBD (n=3 mice per time point). Data represent CBD concentrations as determined by LC/MS for the plasma and brainstem; mean+/-SEM. Blue dashed line represents estimated IC_50_ of CBD for DIPG007 cells. **(H)** PK analysis results for n=3 PPK mice treated with 45 mg/kg CBD. Data represent CBD concentrations as determined by LC/MS for the plasma, normal brain and tumor; mean+/-SEM.

We next assed the pharmacokinetic distribution of CBD in normal brain and brain tumor cells (Fig. 6E-F). After IP administration of a 45 mg/kg dose of CBD, we noted a similar peak concentration of CBD in the brainstem and plasma (6 and 7 uM, respectively) (Fig. 6G), which is above the previously determined IC_50_ dose of CBD in our DIPG cells. At 2 hours, we found equivalent doses of CBD in plasma, brain and brain tumor samples in our PPK model (Fig. 6H).

### CBD treatment in pHGG patients

CBD is increasingly popular as an off-trial, non-prescribed therapy among patients with pHGG (*39*), including DIPG. However, its use remains controversial as no preclinical efficacy, mechanistic data, dosing or clinical studies of CBD in DIPG have been performed. We gathered patient-reported CBD dosing from families of pHGG patients at two institutions (n=15 total, n=8 DIPG, n=11 H3K27M), including patients on an IRB-approved prospective observational study at Children’s Hospital of Colorado for children and young adults with brain tumors undergoing patient-directed medical marijuana therapy (NCT03052738), and retrospective interviews with families of patients who underwent research autopsy at the University of Michigan. CBD was obtained through medical and recreational marijuana dispensaries without prescription; and given orally in all but one case (suppository) one to three times per day with a wide range of dosing (0.07 mg/kg to 25 mg/kg/day, Fig. 7A). No parents reported adverse effects from the CBD aside from taste, and some reported improved nausea and anxiety control. We performed ID1 staining on autopsy samples from high-dose and low-dose H3K27M-mutant tumors. As representative cases, patient UMPED83, with the highest reported dosing (25 mg/kg/day CBD), demonstrated reduced ID1 staining on autopsy sample (Fig. 7B), while UMPED86 underwent low dosing (0.4 mg/kg/day) and demonstrated strong nuclear ID1 staining (Fig. 7C). Patients with pHGG undergoing CBD treatment showed variable ID1 staining in post-mortem tumor tissue, but lower average expression with higher-dose (>3 mg/kg/day) treatments (Fig. 7D).

**Fig. 7.**
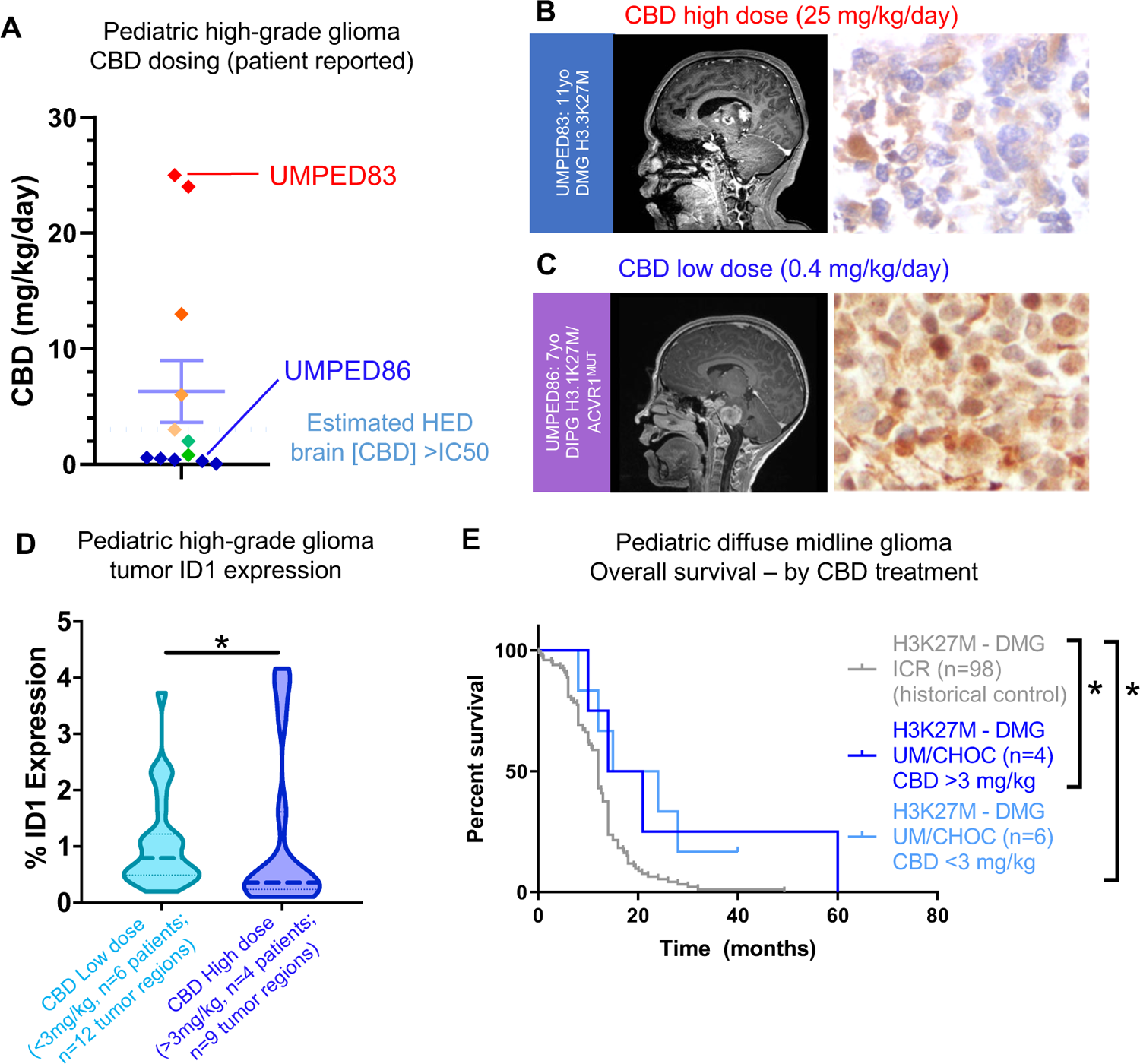
Treatment of human pHGG patients with CBD. **(A)** Plot representing the CBD dosing range (mg/kg/day) in human patients, including one high (UMPED83) and one low (UMPED86) dose of CBD, as indicated by red and blue lines. **(B-C)** IHC-stained tumor tissue from DIPG patients (B) UMPED83 treated with CBD (25 mg/kg) and (C) UMPED86 treated with CBD (0.4 mg/kg/day) during treatment course for assessment of ID1 expression. IHC images representative of n=3 images taken using Aperio ImageScope, magnification=40x. **(D)** ID1 IHC analysis and quantification of human DIPG tumor samples with low (n=6) and high dose (n=4) of CBD; *P=0.0388, Mann-Whitney U test. **(E)** Survival of H3K27M-mutant tumor patients treated with CBD from higher than 3mg/kg (n=4) and lower than <3mg/kg (n=6) with historical control (n=98).

Patients with H3K27M-mutant tumors treated with CBD (n=10) showed improved survival compared to historical controls (*20*), in both high (>3 mg/kg/day) and low (<3 mg/kg/day) CBD treatment groups (Fig. 7E, Supplemental Table 2). These data represent the promise and feasibility of CBD treatment in DIPG, with the clear need for further data in a prospective therapeutic clinical trial.

## DISCUSSION

ID proteins are necessary for appropriate tissue differentiation during embryogenesis, and *ID1* is highly expressed in the normal developing brain followed by quiescence of *ID1* expression in CNS tissue postnatally (*40*). Consistent with the role of ID1 in the pathogenesis of multiple human diseases and cancers (*40–42*), our data indicate that ID1 promotes invasion in DIPG cells, which is a disease-defining feature of this infiltrative tumor. We propose a model by which ID1 is upregulated through multiple mechanisms (H3K27M, ACVR1, region/micro-environment) in order to “re-activate” prenatal brain developmental signaling. Our data support that *ID1^+^* AC-like DIPG tumor cells hijack the transcriptional program of developmental *ID1^+^* OAPC cells in the developing brain cells to produce a “migratory” transcriptional cell state (Fig. 8). We also demonstrate the ability to reverse this ID1-driven phenotype with CBD treatment, and the potential for optimization of this therapeutic targeting.

**Fig. 8.**
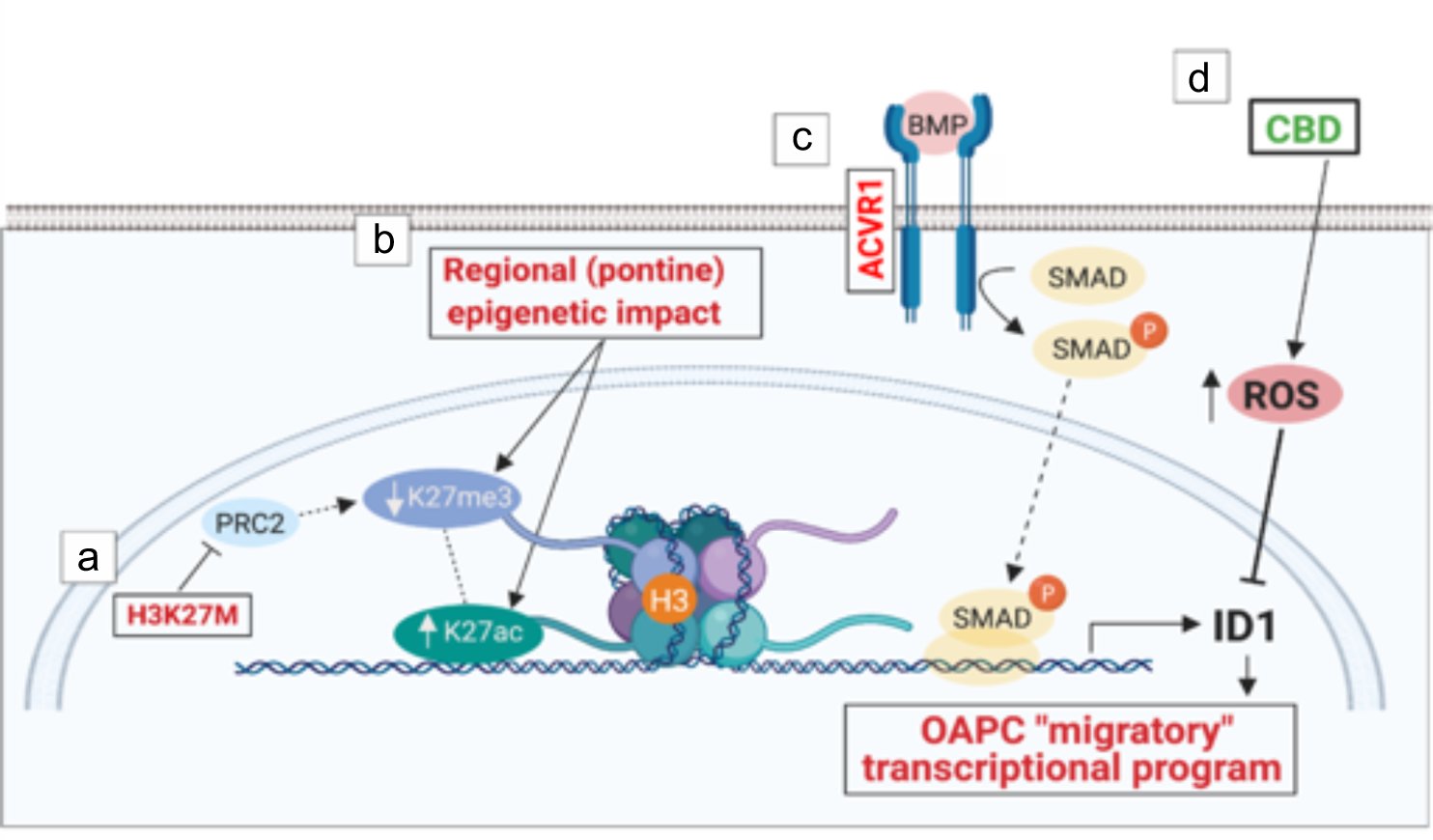
Proposed model of ID1 activation in DIPG with H3K27M and ACVR1 mutations and its inhibition with CBD. The proposed model is made up of the following sub-sections: (a) H3K27M inhibits PRC2, leading to global decreases in H3K27me3 and subsequently allowing for increased H3K27ac. (b) Regional or tissue-specific factors and/ or (c) constitutively activating *ACVR1* mutations increase *ID1* expression via SMAD protein signaling. We propose that ID1 expression replicates the developing cell subtype OAPC transcriptional program, which promotes migration. (d) ID1 expression is reduced by CBD treatment, which partially acts through increasing intracellular levels of reactive oxygen species (ROS). Image created with BioRender.

Our studies implicate an active epigenetic state at the *ID1* locus shared between H3K27M tumor cells and the prenatal precursor brain cells, which is consistent with prior studies focused on H3K27M mutations that have associated changes in H3K27ac/H3K27me3 with differential regulation of key DIPG-associated genes (*43, 44*). Additionally, we provide evidence that post-natal activation of ID1 in tumor cells replicates a prenatal “migratory” transcriptional state seen in a recently discovered subset of developing OAPC brain cells. These OAPCs (*Olig2^+^ SPARCL1^+^HOPX^+^* glial progenitor cells) were recently identified as astrocyte-like at the molecular and transcriptional levels (*28*). In line with this, we found AC-like DIPG cells to transcriptionally mimic the program of OAPCs, with the OAPC-marker *SPARCL1* and *ID1* co-localizing in a subset of H3K27M tumor cells. Interestingly, previous work has suggested a role for *SPARCL1* in promoting DIPG cell invasion into the subventricular zone (SVZ) (*45*).

Secretion of SPARCL1 and pleiotrophin from neural precursor cells (NPCs) was shown to act as a chemoattractant for the DIPG cells, encouraging their infiltration into the SVZ (*45*). Our data demonstrate that *ID1* is most highly expressed by non-cycling AC-like cells in DIPG tumors and *SPARCL1* is one of the strongest expression markers of these cells. This raises the possibility that *SPARCL1* is expressed/secreted *within* DIPG cells, further coordinating or contributing to the invasion of DIPG tumor cells. While further studies are needed to confirm an ID1-driven OAPC/AC-like cellular state, our data raise important insights into the mechanisms underlying one of the most critical and problematic features of DIPG tumors: invasion.

Our data show that ID1 knockdown has the potential to severely impede DIPG tumor cell migration and invasion in pre-clinical models. These phenotypes are consistent with the inherent invasion into normal brainstem tissue that is observed histologically in DIPGs, and with the role of ID1 in other cancers (*11, 42*). In our experiments involving both genetically-engineered and intracranial implantation models, H3K27M-mutant tumors cells with ID1 reduction show reduced tumor growth and invasion.

Cannabidiol is a non-toxic and non-psychoactive member of the endocannabinoid family found in *Cannabis sativa*. CBD has been observed to reduce *ID1* transcription in pre-clinical models of adult cancers (*12*). In the present study, CBD reduced DIPG cell viability and *ID1* expression at concentrations that are likely clinically achievable in the human brain. Our PK studies demonstrated peak brain concentrations of CBD above established IC_50_, despite use of a human equivalent doses (*46*) of only 3 mg/kg, which is well below previously tolerated human CBD dosing. In a phase 1 study, adult patients showed excellent tolerance of oral CBD at 750 mg (15 mg/kg) daily with some non-dose limiting increases in diarrhea and somatic symptoms (muscle ache, fatigue) at 1500 mg (30 mg/kg) daily (*47*). This resulted in peak plasma concentrations of CBD of 1-5 uM (15 mg/kg) and 1.7-10 uM (30 mg/kg) depending on fat content in diet (*47*). Our data showed equivalent plasma and brain concentrations of CBD after IP administration. Previous work has shown that oral administration of CBD in mice results in a 3-4-fold higher concentration in the brain than plasma, likely due to the high lipophilicity of CBD (*35*). CBD is already being used for palliative purposes in pediatric oncology, and CBD has been shown to decrease *ID1* expression and associated oncogenic phenotypes in multiple other cancers in vivo (*11, 12, 48*). Mechanistically, our data suggest that CBD acts to regulate ID1 expression and DIPG cell proliferation partially through increasing intracellular levels of ROS, as previous studies have shown CBD to act through this mechanism in both breast cancer and GBM cells (*37, 38*).

Patients with H3K27M-mutant tumors treated with CBD off-trial show promising improvement in OS compared to historical controls. However, it is important to note that this is limited by the retrospective and heterogeneous nature of our cohort, as well as an unknown number of historical controls that may also have underwent treatment with CBD. Nevertheless, our data make significant strides in establishing the mechanism of this controversial and popular off-trial supplemental compound in high-risk brain tumor patients, and lays the groundwork for future clinical trials. A recent CBD formulation (Epidiolex) has been FDA-approved for epilepsy treatment (*49*), opening the door to a future clinical trial in DIPG (and other ID1-driven tumors).

Our data support a model in which multifactorial genetic and epigenetic processes promote ID1-driven prenatal development transcriptional programs, which also promote the invasive features of DIPG. These results improve our understanding of the pathogenesis of DIPG tumors and provide a strong argument for the inclusion of ID1-targeting therapies into future treatments.

## METHODS

### Study design

The objective of this work was to investigate the role of ID1 in the highly-invasive nature of DIPG and to determine the in vivo antitumor efficacy of genetic and pharmacologic inhibition of ID1 using our IUE *H3.3A*-K27M-mutated murine tumor model. We performed a comprehensive analysis of *ID1* expression by RNA-sequencing of DIPG tissue samples with H3^WT^, H3K27M/ACVR1^WT^, or H3K27M/ACVR^MUT^. We next performed an integrative analysis of H3K27ac and H3K27me3 deposition at the *ID1* gene locus by performing Mint-ChIP-sequencing on these DIPG samples. We further performed transcriptional program analyses of *ID1*-expressing DIPG tumor cells using publicly-available scRNA-seq datasets. To test the in vivo impact of ID1 inhibition, we performed *ID1* knockdown in our PPK tumor model. In vivo pharmacological inhibition of ID1 in our PPK tumor model was performed with CBD. Sample size and any data inclusion/exclusion were defined individually for each mouse experiment. The number of replicates varied between experiments and is presented in figure legends. We performed blinding for quantitative immunohistochemistry scoring of ID1 and Ki67 staining. Finally, we measured ID1 expression in DIPG patient samples which underwent different doses of CBD (non-prescribed) during the course of treatment (Supplement Table 2).

### Murine IUE model of pHGG

All animal studies were conducted according to the guidelines approved by the University Committee on Use and Care of Animals (UCUCA) at the University of Michigan. IUE was performed using sterile technique on isoflurane/oxygen-anesthetized pregnant CD1 females at embryonic stage E13.5, using established methodology. In this study, we injected the following four plasmids together: [1] PBase, [2] PB-CAG-DNp53-Ires-Luciferase (dominant negative TP53 or TP53 hereafter), [3] PB-CAG-PdgfraD824V-Ires-eGFP (PDGFRA D842V), and [4] PB-CAG-H3.3 K27M-Ires-eGFP (H3K27M), referred to as “PPK” model (as previously published) (*17*) (see Supplementary for details).

### Whole exome and transcriptome sequencing (Sick Kids, Toronto)

Use of patient tissues was approved by the Hospital for Sick Children (Toronto) Research Ethics Board. WES/WGS (accession EGAS00001000575) from DIPG samples plus matched normal was using DNA extracted from fresh-frozen tissues as described (*13*). Fresh-frozen tissue was used for total RNA extraction with the RNeasy mini kit (QIAGEN, CA, USA) (see Supplementary for details).

### Mint-ChIP-sequencing

Analyses for the two classical histone modifications H3K27ac and H3K27me3 representing accessible and repressed chromatin states were performed as part of a MiNT-ChIP analysis for 9 tumor samples of DIPG patients in comparison to a control tissue sample of healthy pons according to the protocol published by Buenstro et al., 2013 (see Supplementary for details).

### ScRNA-seq analysis from developing brain and H3K27M-mutant DIPGs

Single-cell gene expression data and their clusters in the developing brain were obtained from GSE133531 (mouse pons), GSE120046 (human pons, gestational week 8-28), and GSE144462 (human cortex, gestational week 21-26) (see Supplementary for details).

### Native ChIP-qPCR

Native ChIP-qPCR was performed on post-mortem tissue using antibodies against H3K27ac (2 µl, cat# 07360, Millipore Sigma), H3K27me3 (1 µg, cat# 07449, Millipore Sigma), and control IgG (2 µg Cat#12370, Millipore Sigma) (see Supplementary for details).

### Invasion assay

Invasion assays were performed using growth factor-reduced Matrigel invasion chambers (Cat #354483, Corning) as previously described (see Supplementary for details) (*50*).

### Migration (scratch) assay

Scratches were made in 80%-confluent 6-well plates, and migration was monitored using the IncuCyte® system (see Supplementary for details).

### CBD treatment studies in murine IUE PPK model

Mice harboring IUE-generated PPK HGG tumors were treated with CBD when tumors reached logarithmic growth phase (minimum 2 x 10^6^ photons/sec via bioluminescent imaging). Mice litters from each experimental group were randomized to treatment with: (A) 15 mg/kg CBD (10% CBD suspended in Ethanol, 80% DPBS, 10% Tween-80) and (B) control treatment (10% Ethanol, 80% DPBS, 10% Tween-80). Mice were treated 5 days/week until morbidity (see Supplementary for details).

### CBD pharmacokinetic analysis

CBD administration to non-tumor bearing CD1 mice and PPK tumor bearing mice for PK studies were performed by IP injection at zero time point. Timeline for CBD injection and plasma, brainstem and/or tumor collection were depicted in Fig. 7 E-F (see Supplementary for details).

### Human studies

Informed consent was obtained for all patient samples. Two patients (CHC001 and CHC002) were enrolled on an ongoing IRB-approved prospective observational study at Children’s Hospital of Colorado for children and young adults with brain tumors undergoing patient-directed medical marijuana therapy (NCT03052738) (see Supplementary for details).

## Acknowledgments

The authors thank the patients and their families for participation in this study.

## Author contributions

Conception and design of study: VNY, MKH, CK Acquisition, analysis, or interpretation of data: VNY, MKH, DM, CT, JRC, TY, RW, RoS, MB, SS, TQ, BM, RuS, RR, MN, KFG, MAHG, KD, NAV, JL, MGF, XC, MGC, PRL, RM, AC, PYD, SM, CH, SMW, SV, CK Drafting and revising the written manuscript: VNY, MKH, JRC, RuS, CK Final approval of version to be published: VNY, MKH, DM, CT, JRC, TY, RW, RoS, MB, SS, TQ, BM, RuS, RR, MN, KFG, MAHG, KD, NAV, SVS, JS, JL, MGF, XC, MGC, PRL, RM, AC, PYD, SM, CH, SMW, SV, CK

## Data and materials availability

All data are available in the main text or the supplementary materials.

## Supplementary Materials

### Supplementary Methods

#### Whole exome and transcriptome sequencing (Sick Kids, Toronto)

Use of patient tissues was approved by the Hospital for Sick Children (Toronto) Research Ethics Board. WES/WGS (accession EGAS00001000575) from DIPG samples plus matched normal was using DNA extracted from fresh-frozen tissues as described (*13*). Fresh-frozen tissue was used for total RNA extraction with the RNeasy mini kit (QIAGEN, CA, USA). 34 DIPG and 17 normal brain samples passed quality control. The TruSeq Stranded Total RNA Library Prep with Ribo-Zero Gold Kit (Illumina, CA, USA) was used and paired end sequencing generated with Illumina HiSeq 2500 machines (accession EGAD00001006450) (*51*). Sequencing quality was confirmed with FastQC v0.11 (http://www.bioinformatics.babraham.ac.uk/projects/fastqc/). Reads were quality trimmed with Trimmomatic (*52*) v0.35 before being aligned with RSEM (*53*) v1.2 to human transcriptome build GRCh37 v75. Gene expression was quantified FPKM.

#### Whole exome and transcriptome sequencing of tumor/normal tissue (University of Michigan)

Clinically integrated sequencing was performed according to previously published methodology (*54, 55*). For living patients with DIPG/HGG, the PEDS-MIONCOSEQ study was approved by the Institutional Review Board of the University of Michigan Medical School and all patients or their parents or legal guardians provided informed consent (written assent if >10 years). For deceased patients, parents were consent for research autopsy and brain tumor/normal banking separately from the MIONCOSEQ protocol. Tumor (FFPE or frozen) and normal (cheek swab or blood, when available) samples were submitted for whole exome (paired tumor and germline DNA) and transcriptome (tumor RNA) sequencing. Nucleic acid preparation, high-throughput sequencing, and computational analysis were performed by the Michigan Center for Translational Pathology (MCTP) sequencing laboratory using standard protocols in adherence to the Clinical Laboratory Improvement Amendments (CLIA) (*56*).

#### Analysis of tumors from Institute for Cancer Research (ICR)

Whole exome and transcriptome sequencing data from 1067 pediatric high grade gliomas (pHGGs) (compiled from the Jones lab, ICR London, Cancer Cell 2017) was retrieved from the ICR cohort (*20*). Specimens with mRNA sequencing (n=247) were then separated by location into brainstem/pons (n=68), hemispheric (n=130), and midline (n=49). PHGGs of the brainstem were considered DIPGs. Of the 68 DIPGs from the ICR cohort, 2 did not have survival data and were removed. Overall survival was defined from day of diagnosis to death of patient. High *ID1* expression was defined as having a z-score greater than 0.2 (n=38), and low *ID1* expression as less than 0.2 (n=25).

#### In Utero Electroporation (IUE) and generation of primary cell lines from IUE tumors

PiggyBac transposon plasmids containing *PDGFRA* mutation, *TP53* mutation, *H3F3A-* K27M, and *H3F3A-*WT, were kind gifts from Dr. Timothy Phoenix (Cincinnati Children’s Hospital, Cincinnati, OH) (*57*). *In utero* electroporation was performed on isoflurane/oxygen-anesthetized pregnant female mice at embryonic day E13.5 in the cortex. Subcutaneous delivery of Vetergesic and Carpofen at 0.1 mg/kg and 5 mg/kg, respectively, was also provided pre-emptively. Briefly, IUE were performed using sterile technique on isoflurane/oxygen-anesthetized pregnant CD1 females at E13.5. Uterine horns were exposed through a 1 cm incision and embryos were digitally manipulated into the correct orientation. Borosilicate capillaries were loaded with endotoxin-free DNA and Fast Green dye (0.05%, Sigma) for visualization. Lateral ventricles were then injected with the DNA-dye mixture using a microinjector (Eppendorf). 3-5 plasmids were injected at the same time, each at a concentration of 2 µl/µl. 1-2 µl of total solution was injected into each embryo. DNA was electroporated into cortical neural progenitors using 3 mm tweezertrodes (BTX), applying 5 square pulses at 35 V, 50 ms each with 950 ms intervals. Embryos were then returned into abdominal cavity, muscle and skin sutured, and animal monitored until full recovery. Periodically, tumor growth was monitored by IVIS as mice are treated starting 33 days post injection (dpi).

Primary cell lines with specific genetic alterations were generated from IUE-induced pediatric high grade glioma models. Mice with confirmed large tumors (bioluminescence 10^7^ – 10^8^ photons/s/cm^2^/sr) were selected. Mice were euthanized with an overdose of isoflurane, decapitated, and brain was dissected from the skull. Brain was then placed in a Petri dish, and coronal cuts were made anterior and posterior to tumor using sterile scalpel. Tumor was identified and dissected with fine forceps and placed in a 1.5 ml tube containing 300 µl of Neural Stem Cell Media (NSC Media: DMEM/F12 with B-27 supplement, N2 supplement, and Normocin, supplemented with human recombinant EGF and bFGF at a concentration of 20 ng/ml each). Tumor was gently homogenized using a plastic pestle. 1 ml of enzyme free tissue dissociation solution was added to homogenized tumor, and then incubated at 37°C for 5 minutes. Then, cell suspension was passed through a 70 µm cell strainer, centrifuged at 300x g for 4 min. Supernatant was decanted, and pellet resuspended in 7 ml of NSC media. Solution was then plated onto a T25 tissue culture flask, and placed in tissue culture incubator at 37°C with atmosphere of 95% air and 5% CO_2_. After 3 days, neurospheres were removed and re-plated into a T75 tissue culture flask. Cells were then maintained in NSC media appropriately. No mycoplasma testing regimen was performed on murine cell lines as they are early passage tumor-derived cells. If frozen, cells were cultured for 2 to 3 passages (2 weeks) following thawing for experiments.

#### Mint-ChIP-sequencing of tumor tissue

Analyses for the two classical histone modifications H3K27ac and H3K27me3 representing accessible and repressed chromatin states were performed as part of a MiNT-ChIP analysis for 9 tumor samples of DIPG patients in comparison to a control tissue sample of healthy pons according to the protocol published by Buenstro et al., 2013. Up to 50 mm³ snap frozen tumor tissue was digested with 2.5 mg/ml collagenase IV (Sigma-Aldrich, Germany) and dissociated via the gentleMACS Dissociator (Miltenyi, Germany). Subsequent immunoprecipitation for H3K27Ac and H3K27me3 was performed with 5 µg of ChIP-grade antibodies, monoclonal murine anti-H3K27Ac (MABI0309, ActiveMotif, Belgium) and a polyclonal rabbit anti-H3K27m3 (Merck Millipore, Germany).

Over 50 mio reads were sequenced in 50 bp paired-end sequencing runs on a NovaSeq 6000 system (NGS Core Facility, University Hospital, Bonn, Germany) and demultiplexed as described by Buenstro et al., 2013 (Core Unit Bioinformatics Data Analysis, University Hospital Bonn, Germany). Reads were aligned against the human reference genome hg19 by Bowtie2 (v2.4.2). Tag directories of piled up reads were created using HOMER (v4.11) makeTagDirectory and visualized makeUCSCfile with the -fsize 5e8 option.

#### Native ChIP-qPCR

Protocol for native ChIP-qPCR was adapted from previously described methods, and optimized for frozen human tissue (*58*). Antibodies against H3K27ac (2 µl, cat# 07360, Millipore Sigma), H3K27me3 (1 µg, cat# 07449, Millipore Sigma), and control IgG (2 µg Cat#12370, Millipore Sigma) were used for immunoprecipitation.

Quantitative-PCR was performed per below methods, using 1µl of eluted ChIP DNA. Primers for *ID1* enhancer and promoter region target sites were predicted based on H3K27ac peaks observed in the four H3K27M DIPG tumor tissue samples analyzed via ChIP-sequencing in main Figure 2C. For a complete list of primers used in ChIP-qPCR, see Supplementary Table S1. NCBI RefSeq hg19 was used as reference genome (*43*). Enrichment at target sites was quantified using the percent input method as has been previously described (*59*). Gene expression was quantified relative to GAPDH using the comparative C_T_ method as previously described (*60*). For a complete list of primer sequences used in qPCR for gene expression, see Supplementary Table S2.

#### Analysis of developing murine brain

Call sets from the ENCODE portal (https://encodeproject.org/) were downloaded with the following identifiers: ENCSR691NQH, ENCSR428GHF, and ENCSR066XFL. ChIP-Sequencing peaks were quantified using EaSeq (http://easeq.net) (*61*). Graphic depictions of H3K27ac peaks at the *ID1* locus were generated using IGV browser (*62*). ID1 in situ hybridization (ISH) data and images from the 2014 Allen Developing Mouse brain Atlas (http://developingmouse.brain-map.org/) were downloaded and analyzed.

#### ScRNA-seq analysis from developing brain and H3K27M-mutant DIPGs

Single-cell gene expression data and their clusters in the developing brain were obtained from GSE133531 (mouse pons), GSE120046 (human pons, gestational week 8-28), and GSE144462 (human cortex, gestational week 21-26). Raw mouse expression data was normalized to counts-per-million for each cell. Cells were assigned to clusters based on the joint clustering of cells from all four developmental stages (E15.5, P0, P3, P6). 1,792 cells were removed due to missing cluster assignments and Id1 expression was analyzed in the remaining 22,682 cells. Analysis of normalized human pontine expression data was restricted to 4,228 cells that were detected across 18 gestational time points in the pons (>=3 cells per gestational week). Normalized human expression data for H3K27M-mutant DMGs was obtained from GSE102130. Tumor cells with an astrocytic differentiation (AC-like), oligodendrocytic differentiation (OC-like), and OPC-like program were determined using stemness- and lineage scores from Filbin et al. (*19*) and *k*-means clustering. Mann-Whitney U (MWU) tests were used to identify for each patient genes that separate AC-like and OPC-like cells. Cell type enrichments were calculated using significant marker genes (cell type set A) and full summary statistics obtained from differential marker gene analysis (enrichment score=z-transformed median -log10 MWU P values). Functional enrichment analysis of marker genes was performed using the Enrichr web service (*63*) and top 200 marker genes (sorted by MWU P-value).

### DIPG immunohistochemistry (IHC) staining and quantification

Mouse PPK tumor and human DIPG paraffin embedded tissue were sectioned and sent to Dr. Daniel Martinez (Department of Pathology, Children’s Hospital of Philadelphia, PA) for ID1 and Ki67 staining. Briefly, ID1 antibody (Biocheck BCH-1) was used to stain formalin-fixed paraffin embedded tissue slides. Slides were rinsed in 2 changes of xylene for 5 min each then rehydrated in a series of descending concentrations of ethanol. Slides were treated with .3% H2O2/methanol for 30min. and then treated in a pressure cooker (Biocare Medical) with 0.01M Citrate buffer pH 7.6. After cooling, slides were rinsed in 0.1M Tris Buffer and then blocked with 2% fetal bovine serum for 5 min. Slides were then incubated with ID1 antibody at a 1:25 dilution overnight at 4 degrees C. Slides were then rinsed and incubated with biotinylated anti-Rabbit IgG (Vector Laboratories BA-1000) for 30min at room temp. After rinsing, slides were incubated with the avidin biotin complex (Vector Laboratories PK-6100) for 30 min at room temp. Slides were then rinsed and incubated with DAB (DAKO Cytomation K3468) for 10 min at room temp. Slides were counterstained with hematoxylin, then rinsed, dehydrated through a series of ascending concentrations of ethanol and xylene, then coverslipped. Ki67(SP6) antibody (Abcam ab16667) was used to stain formalin-fixed, paraffin–embedded tissue. Staining was performed on a Bond Max automated staining system (Leica Microsystems). The Bond Refine staining kit (Leica Microsystems DS9800) was used. The standard protocol was followed with the exception of the primary antibody incubation which was extended to 1 hour at room temperature. Ki67 was used at 1:400. Antigen retrieval was performed with E2 (Leica Microsystems) retrieval solution for 20min. After drying, slides were scanned at 20x magnification with an Aperio CS-O (Leica Biosystems) slide scanner and images were viewed using the Aperio ImageScope software. An individual blinded to the experiment captured five random images from each IHC slide at 10X magnification. Quantification of images for precent positive area were measured by ImageJ software.

### Human cell cultures

Primary H3.3K27M-mutant cell line DIPG007 was obtained from Dr. Rintaru Hashizume from (Northwestern University, Chicago, IL) who obtained them originally from Dr. Angel Carcaboso (Hospital Sant Joan dr Deu, Barcelona, Spain). DIPG-XIII was obtained from Dr. Michelle Monje (Stanford University, Stanford, CA). PBT-29 was obtained from Dr. Nicholas Vitanza,(Seattle Children’s, Seattle, WA). Immortalized human embryonic kidney 293 (HEK293) cells were obtained from Dr. Sriram Venneti (University of Michigan, Ann Arbor, MI). Cells were cultured for 2 to 3 passages (2 weeks) following thawing for experiments.

#### DIGP007, DIPGXIIIp and PBT-29 cells

DIPG007, DIPGXIIIp and PBT-29 cells were cultured in TSM N5 media: 250 ml DMEM (1X, Cat#11995065, Gibco); 250 ml NeuroBasal-A Medium (1X, Cat#0888022, Gibco); 5 ml HEPES (1M, Cat#15630080, Gibco); 5 ml Sodium Pyruvate (100mM, Cat#11360070, Gibco); B-27 Supplement without Vitamin A (50X, Cat #12587010, Gibco); 5 ml MEM NEAA (100X, Cat#11140050, Gibco); 5 ml Antibiotic-Antimycotic (100X, Cat#15240062, Gibco); 250 µl Heparin Solution (Cat#07980, STEMCELL Technologies); 10 µl human PDGF-AA every 3 days (10 ng/ml, Cat#10016, Shenandoah Biotechnology); 1 ml Normocin (Cat#antnr1, InvivoGen); 10µl human PDGF-BB every 3 days (10 ng/ml, Cat#10018, Shenandoah Biotechnology); 20 µl FGF every 3 days (20 ng/ml, Cat#10018B, PeproTech); 20 µl EGF every 3 days (20 ng/ml, Cat#10047, PeproTech). For adherent conditions, FBS was diluted in media to 10%. For neurosphere culture, FBS was not added. At each passage, cells were dissociated using StemPro Accutase (Cat#A1110501, Gibco).

#### Human Embryonic Kidney 293 (HEK293) cells

HEK293 cells were cultured in: 500 ml DMEM (1X, Cat#11995065, Gibco); 333 µl Gluta-Max (200 mM, Cat#25030081, Gibco); 1 ml Normocin (Cat#antnr1, InvivoGen). FBS was diluted in media to 10%. At each passage, cells were dissociated using StemPro Accutase (Cat#A1110501, Gibco).

### ShRNA-mediated gene silencing by lentiviral transduction of cultured cells

ShRNA-mediated gene silencing for DIPG007, HEK293, or NHA cell cultures was performed by lentiviral transduction with pGIPZ shRNAs (Dharmacon, GE) targeting *ID1* (Clone ID’s V2LHS_133263, V2LHS_133264) or scrambled control (Cat#RHS4346). A map of this vector is provided in Supplementary Figure S15. Protocol for lentiviral transduction was modified from the University of Michigan Vector Core as follows. 24 hours prior to transduction, cells were split into 6-well tissue culture plates at a density that they would reach approximately 60% confluency the following day. The next day, media was aspirated and replaced with 1.35 ml of fresh media. Then, 0.15 ml of 10x viral supernatant was added, along with 2.5 µl of 4mg/ml Polybrene (Cat#G062, ABM). Plate was then rocked gently on shaker to evenly distribute virus and Polybrene. Cells were then placed in cell incubator at 37°C for approximately 24 hours. Exact time was dependent on when cells began expressing GFP, which was contained in the lenti-vector.

### Western Blotting

Western blotting was performed using antibodies against ID1 (1:1000, Cat#133104, Santa Cruz Biotechnology), Vinculin (1:10000, Cat#700062, Invitrogen), H3K27M (1:500, EMD, Cat#ABE419), H3K27me3 (1:500, EMD, Cat#07-449) and ACTB (1:10000, Cat#A2228, Sigma-Aldrich), Secondary antibodies biotinylated horse anti-mouse IgG (Cat#BA2000, Vector Laboratories), HRP goat anti-rabbit IgG (Cat#PI1000, Vector Laboratories), and m-IgG_k_ BP-HRP (Cat#sc516102, Santa Cruz Biotechnology) were used. Chemiluminescent blots were imaged and processed using the FluroChem M system (ProteinSimple, San Jose, CA).

### Cannabidiol treatment studies in vitro

Treatment was performed as previously described (*12*). 3,000 primary DIPG007 and PPK cells were plated in 96-well plates and incubated for 24 hours. The next day, cells were treated with different doses of cannabidiol (CBD) cat # 90080 (Cayman Chemical). After 72 hours, in vitro cell viability was monitored by XTT Cell Proliferation Assay kit (Cayman Chemical).

### Invasion assay

Invasion assays were performed using growth factor-reduced matrigel invasion chambers with 8 uM pores (Cat #354483, Corning) as described in previously published work (*50*). Seeding density and incubation time was optimized for each cell line. FBS was used as chemoattractant. Invading cells were stained with crystal violet. To count invading cells, transwell membranes were viewed underneath an inverted microscope at 10x magnification, and four pictures were taken at random locations to get an average sum.

### Migration (scratch) assay

Migration assays were performed following a previously published protocol with slight modifications (*64*). Cells were seeded in 6-well plates, and grown to approximately 80% confluence. Scratches were made using a 200 µl pipette tip, and migration was then monitored using the IncuCyte® live-cell analysis system (Sartorius, Ann Arbor, MI). Images were analyzed using ImageJ’s MRI Wound healing tool (http://dev.mri.cnrs.fr/projects/imagej-macros/wiki/Wound_Healing_Tool). Percent closure was calculated as [(*Area_t=0_* – *Area_t_) /Area_t=0_*]*100.

### Proliferation and viability assays

Cell viability was quantified using the MTT Cell Proliferation Assay Kit (Cat#ab211091, ABCAM), following manufacturer instruction for adherent cells. For proliferation, cells were seeded in 96-well plates and monitored for confluence using the IncuCyte® live-cell analysis system (Sartorius, Ann Arbor, MI).

### Implantation of DIPG007 cells and bioluminescence imaging

#### Implantation of mouse cells

Male and female NSG™ mice were obtained from Jackson Labs (Bar Harbor, ME) and were 6-10 weeks of age at the start of surgery. All animal studies were conducted according to the guidelines approved by the Institutional Animal Care & Use Committee (IACUC) at the University of Michigan. Mice were anesthetized with injection of 120 mg/kg ketamine and 0.5 mg/kg dexmedetomidine. Hair above scalp was shaven, disinfected with iodine, and a 1 cm incision was made above scalp to expose cranium. The periosteum was removed with scalpel. Next, a 0.6mm burr hole was drilled 2 mm right of midline and 0.2 mm anterior to the bregma with the Ideal Micro Drill (MD-1200 120V) from Braintree Scientific Inc. Mice were placed in a Mouse/Neonatal Rat Adaptor stereotactic frame (#51615) from Stoelting. A 10 ul syringe (#7635-01) fitted with 33-gauge needle (#7762-06) from Hamilton, was filled with cell suspension (15,000 cells per uL) and penetrated 3 mm into brain tissue. After waiting two minutes, one microliter of cell suspension was injected over one minute and needle was slowly removed after waiting 3 minutes after injection. Incision was closed with 4-0 nylon and mouse was given 1 mg/kg atipamezole for reversal and monitored for recovery. Mice were monitored for symptoms of morbidity, including impaired mobility, scruffed fur, hunched posture, ataxia, and seizures.

#### Bioluminescence imaging

Mice were imaged using IVIS Spectrum #2 machine at the Center for Molecular Imaging at the University of Michigan Core Facility. Mice were injected with 160 mg/kg D-luciferin (#115144-35-9) from Gold Biotechnology and anesthetized with 2% isoflurane. 10 minutes after luciferin injection, mice were placed into machine in a prone position and bioluminescence was measured. Mice were imaged until peak signal was obtained for each mouse. Tumor bioluminescent signal is measured in radiance (photons) (p/s/cm^3^/sr) in a circular region of interest (ROI) over the cranium of each mouse with Living Image Software (PerkinElmer Inc).

### CBD treatment studies in murine IUE PPK model

Mice harboring IUE-generated PPK HGG tumors were treated with CBD when tumors reached logarithmic growth phase (minimum 2 x 10^6^ photons/sec via bioluminescent imaging). Mice litters from each experimental group were randomized to treatment with: (A) 15 mg/kg CBD (10% CBD suspended in Ethanol, 80% DPBS, 10% Tween-80) and (B) control treatment (10% Ethanol, 80% DPBS, 10% Tween-80). Mice were treated 5 days/week until morbidity.

Animals displaying symptoms of morbidity after treatment were euthanized for immunohistochemistry (IHC) analysis. For IHC analysis, mice were perfused with Tyrode’s Solution followed by 4% paraformaldehyde fixative solution to preserve the structures of the brain. For IHC quantification (Ki67 and ID1), 3-4 random images per tumor (n=3 tumors per group) were taken at 10x magnification using Aperio ImageScope and percent positive area was calculated using ImageJ software.

### CBD Pharmacokinetic analysis

#### Mouse PK sample procurement

CBD administration to non-tumor bearing CD1 mice and PPK tumor bearing mice for PK studies were performed by IP injection at zero time point. Timeline for CBD injection and plasma, brainstem and/or tumor collection were depicted in Fig. 7 E-F. At half, one, two, and six hours after the CBD injection, the mice were isoflurane/oxygen-anesthetized and 500 uL to 1 mL of blood was drawn from the apex of the heart within the mouse’s enclosed cavity. Immediately, the withdrawn blood was centrifuged within a microvette EDTA coated conical tube for 10 minutes at 10,000 RPM, and the plasma was separated and stored at −80°C until PK analysis was performed. Following the blood draw, the mouse was sacrificed and the brain, brain stem, and/or tumor were extracted separately and stored at −80°C until PK analysis was performed.

#### Chemicals and reagents

For PK studies, CBD powder was procured from Cayman chemical USA. Liquid chromatography–mass spectrometry (LC-MS) grade acetonitrile was purchased from Sigma-Aldrich. Formic acid (98%; LC-MS grade) was obtained from Fluka. A Milli-Q water system from Millipore was used to obtain ultrapure deionized water.

#### Sample preparation

Plasma (40 µL) was dispensed into a Fisher Scientific 96-well plate, to which 40 µL of ice-cold acetonitrile (100%) and 120 µL of internal standard solution (1000 ng/mL) were added. Next, the plate was vortexed for 10 minutes. The plate was then centrifuged at 3500 revolutions per minute (RPM) for 10 minutes at 4°C to precipitate the protein. LC–tandem mass spectrometry (LC-MS/MS) was used to analyze 5 µL of the supernatant. The plasma samples were sonicated prior to being transferred to the 96-well plates. Tissue samples were weighed and suspended in 20% acetonitrile (80% water; 1:5 wt/vol). The samples were then homogenized four times for 20 seconds each time at 6,500 RPM in a Precellys Evolution system. For LC-MS/MS analysis, the CBD in brain tissue homogenates were extracted from the samples in the same manner as the CBD in plasma. Prior to extraction, samples that were above the upper limit of qualification were diluted with the same matrix. Calibrator-standard samples and quality control samples were prepared by mixing 40 µL of blank bio matrix, 40 µL of working solution, and 120 µL of internal standard solution.

#### Calibration curve

Analytical curves were made with 12 nonzero standards by plotting the peak area ratio of CBD to the internal standard vs the concentration. The curve was created with linear regression and weighted (1/X2). The correlation coefficient demonstrated the linearity of the relationship between peak area ratio and concentration.

#### Liquid chromatography tandem–mass spectrometry

The concentrations of CBD were determined with a Sciex AB-5500 Qtrap mass spectrometer with electrospray ionization source, interfaced with a Shimadzu high-performance LC system. The LC-MS/MS system was controlled with Analyst Software version 1.6 from Applied Biosystems; this was also used for acquisition and processing of data. Separation was performed on a Waters Xbridge C18 column (50 × 2.1 mm ID, 3.5 µm); the flow rate was 0.4 mL/min. A (100% H2O with 0.1% formic acid) and B (100% acetonitrile with 0.1% formic acid) comprised the mobile phase. The gradient began with 5% B for 30 seconds and then linearly increased to 99% B at 2 minute and then reduced to 5% B at 4.1 minutes to 5.5 minutes with a runtime of 6 minutes in total. The mass spectrometer was operated in positive mode; multiple reaction monitoring was used for analysis. The Q1 m/z and Q3 m/z was 487.9 and 401.1, respectively.

### Statistical analyses

Statistical analyses were performed in consultation with a bioinformatician. Graphs were plotted and statistical analyses were performed using GraphPad Prism software (version 7.00/8.00, GraphPad, La Jolla, CA) and Microsoft Excel. Unpaired, two-sided analysis of variance (ANOVA) followed by multiple comparison analyses were used to analyze data as indicated. Survival analyses in animals were performed using Kaplan-Meier analyses with the Log-Rank test. Data were considered significant if p values were below 0.05 (95% confidence intervals).

### Human studies

Informed consent was obtained for all patient samples. Two patients (CHC001 and CHC002) were enrolled on an ongoing IRB-approved prospective observational study at Children’s Hospital of Colorado for children and young adults with brain tumors undergoing patient-directed medical marijuana therapy (NCT03052738). The University of Michigan cohort consisted of retrospective interviews with families of patients who underwent research autopsy. The patients all underwent research autopsy consent and were contacted to confirm use of patient details and tumor samples for this study. Patients who reported CBD therapy at any point in their care were included in this study, and CBD dosage was confirmed by pictures of CBD bottle, discussion with dispensary, etc., when possible.

## Supplementary figures

**Supplementary Figure S1.**
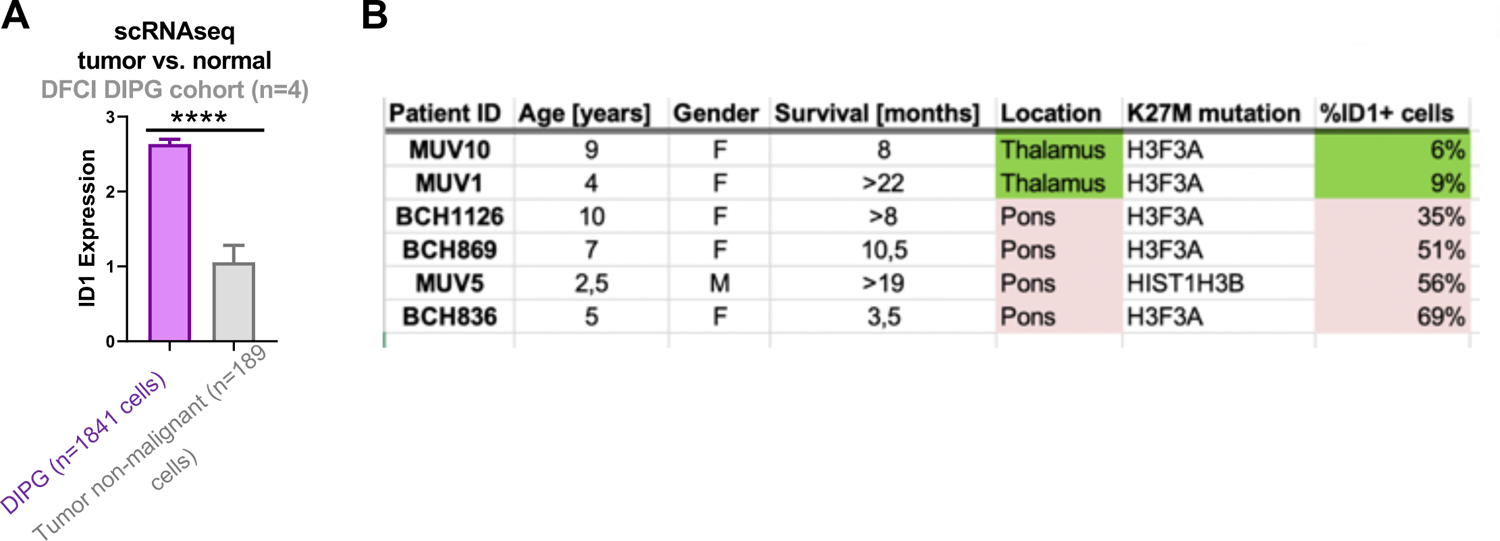
*ID1* expression in DIPG by cell malignancy and tumor location. **(A)** *ID1* expression of DIPG tumor by cell malignancy from the Dana-Farber Cancer Institute (DFCI) DIPG cohort (n=4 patients). ID1 expression was compared between malignant DIPG cells (n=1841) and non-malignant tumor cells (n=189) from single-cell RNA-seq (scRNA-seq) data. Data represent mean +/- SEM; ****P<0.0001, unpaired parametric t test. **(B)** *ID1* is frequently (35-69%) expressed in pontine DIPG cells and rarely (6-9%) expressed in thalamic DMGs.

**Supplementary Figure S2.**
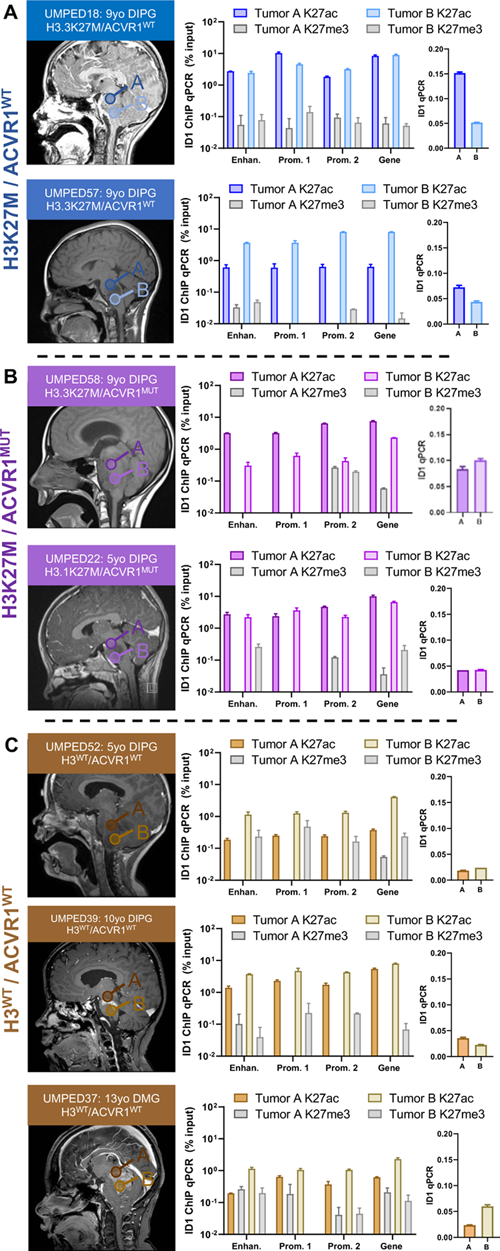
Multifocal ChIP-qPCR analysis of *ID1* expression in human DIPG. **(A-C)** [Left panel]: Multifocal DIPG tumor samples (2 per tumor) were obtained at autopsy from n=2 patients with H3K27M mutation and wildtype *ACVR1* (ACVR1^WT^), n=2 patients with H3K27M mutation and *ACVR1* mutation (ACVR1^MUT^) and n=3 patients with wildtype H3 (H3^WT^) and *ACVR1*. Circles labeled “A” and “B” over MRI images represent the approximate region of tumor where a sample was obtained from. [Right panel]: Graphs on left represent percent relative enrichment for H3K27ac and H3K27me3 marks by ChIP-qPCR for each of the predicted *ID1* gene body elements shown in main figure 2. Graphs on right represent *ID1* expression, measured by qPCR, for the multifocal samples collected from patients in shown in left MRI images. Data represent mean +/- SEM.

**Supplementary Figure S3.**
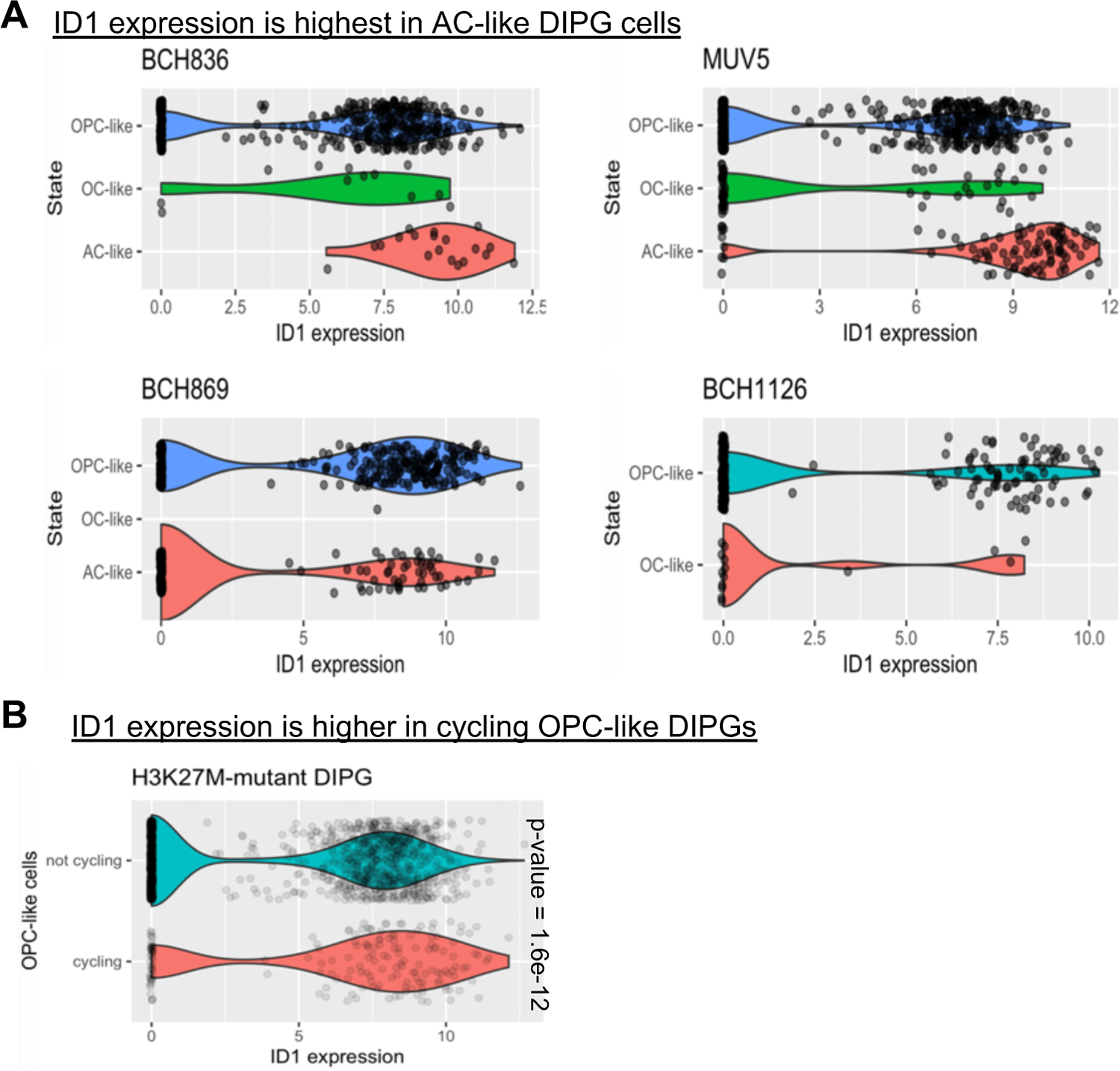
*ID1* expression from single cell RNA-sequencing of six different H3K27M-DMG patients across varying regions and malignant cell types. **(A)** Violin plots depicting ID1 expression in three subtypes of H3K27M-DIPG malignant cells [Data from pontine DIPG patients in Fig. 1B]. (B) Violin plots depicting ID1 expression in cycling vs non-cycling malignant H3K27M-DIPG cells; P=1.6e^-12^, Mann-Whitney U test. [OPC-Oligodendrocyte precursor cell; OC-Oligodendrocyte; AC-Astrocyte]. Primary data for parts (A) and (B) from Filbin et al., *Science*, 2018.

**Supplementary Figure S4.**
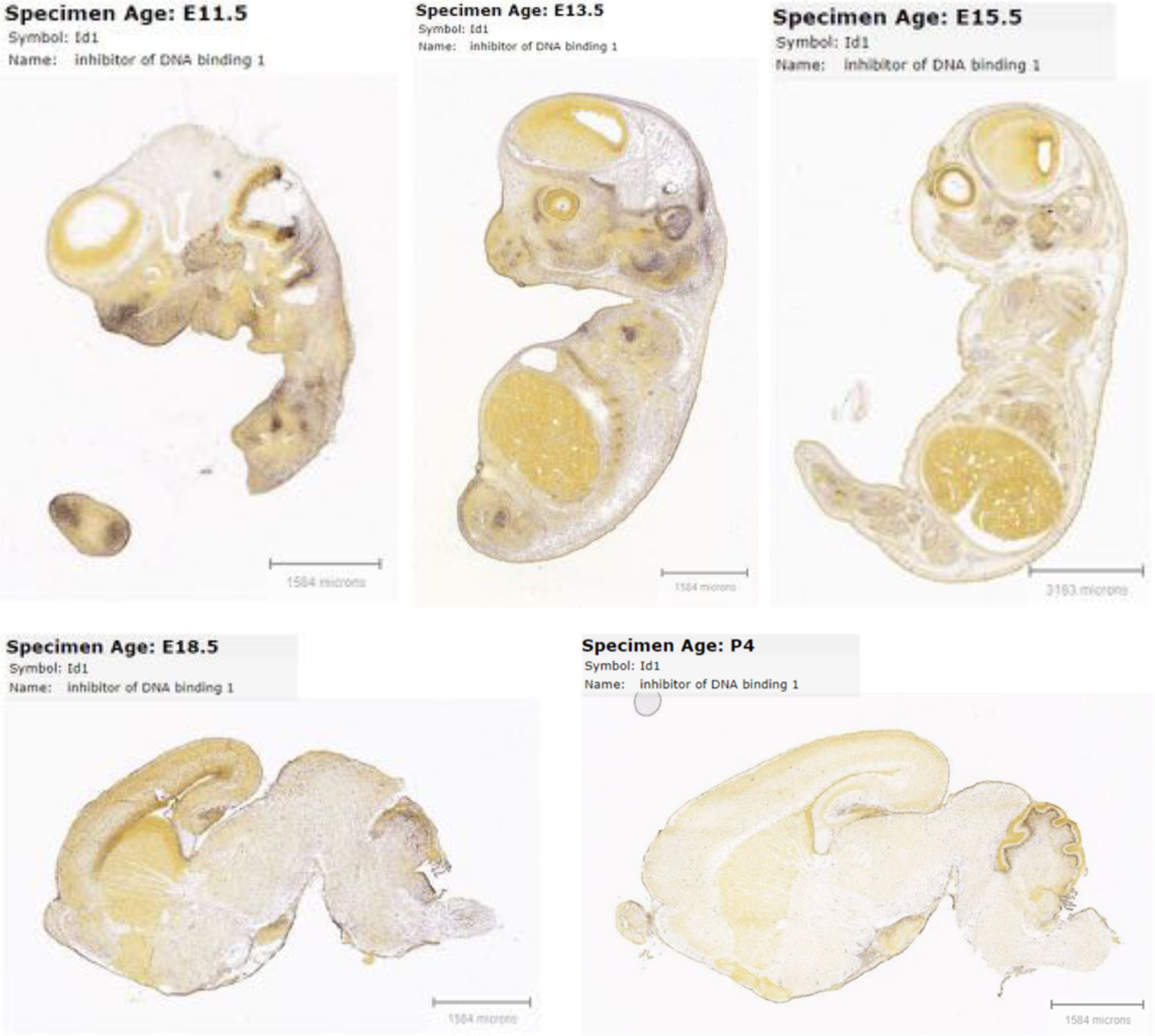
In situ hybridization for *ID1* RNA in developing mouse brain. *ID1* RNA is high in the developing embryonic murine brain, and drastically reduced in the post-natal brain. Image credit: Allen Institute. © 2014 Allen Developing Mouse Brain Atlas. Available from: http://developingmouse.brain-map.org/

**Supplementary Figure S5.**
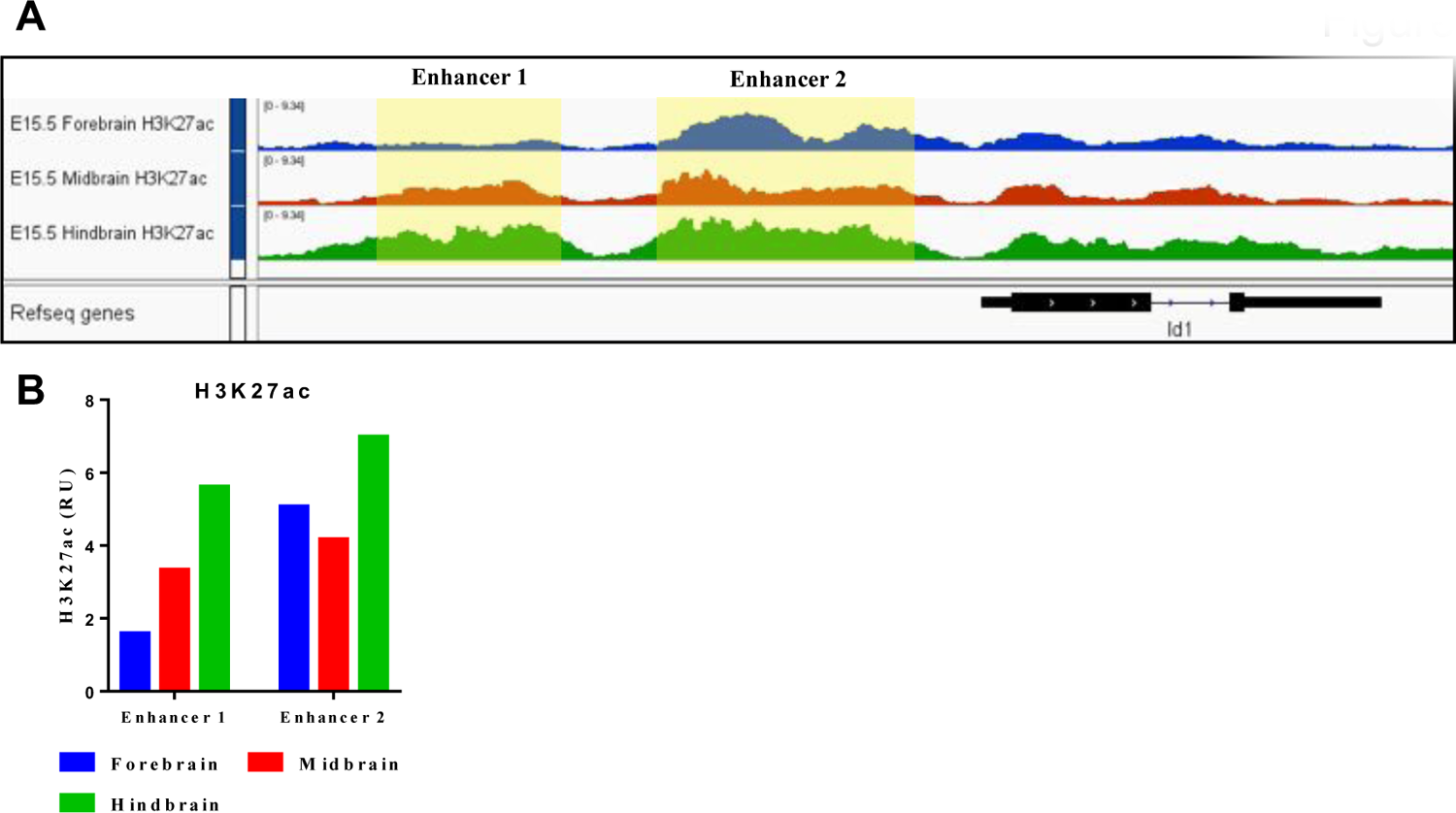
H3K27ac at *ID1* locus during murine development. **(A)** H3K27ac peaks at the ID1 locus in E15.5 mouse brain and predicted ID1 enhancer regions [Image generated using IGV browser]. **(B)** Relative enrichment of H3K27ac at predicted ID1 enhancers in E15.5 murine brain regions. Data retrieved from ENCODE Consortium; highlighted regions quantified using EaSeq (http://easeq.net).

**Supplementary Figure S6.**
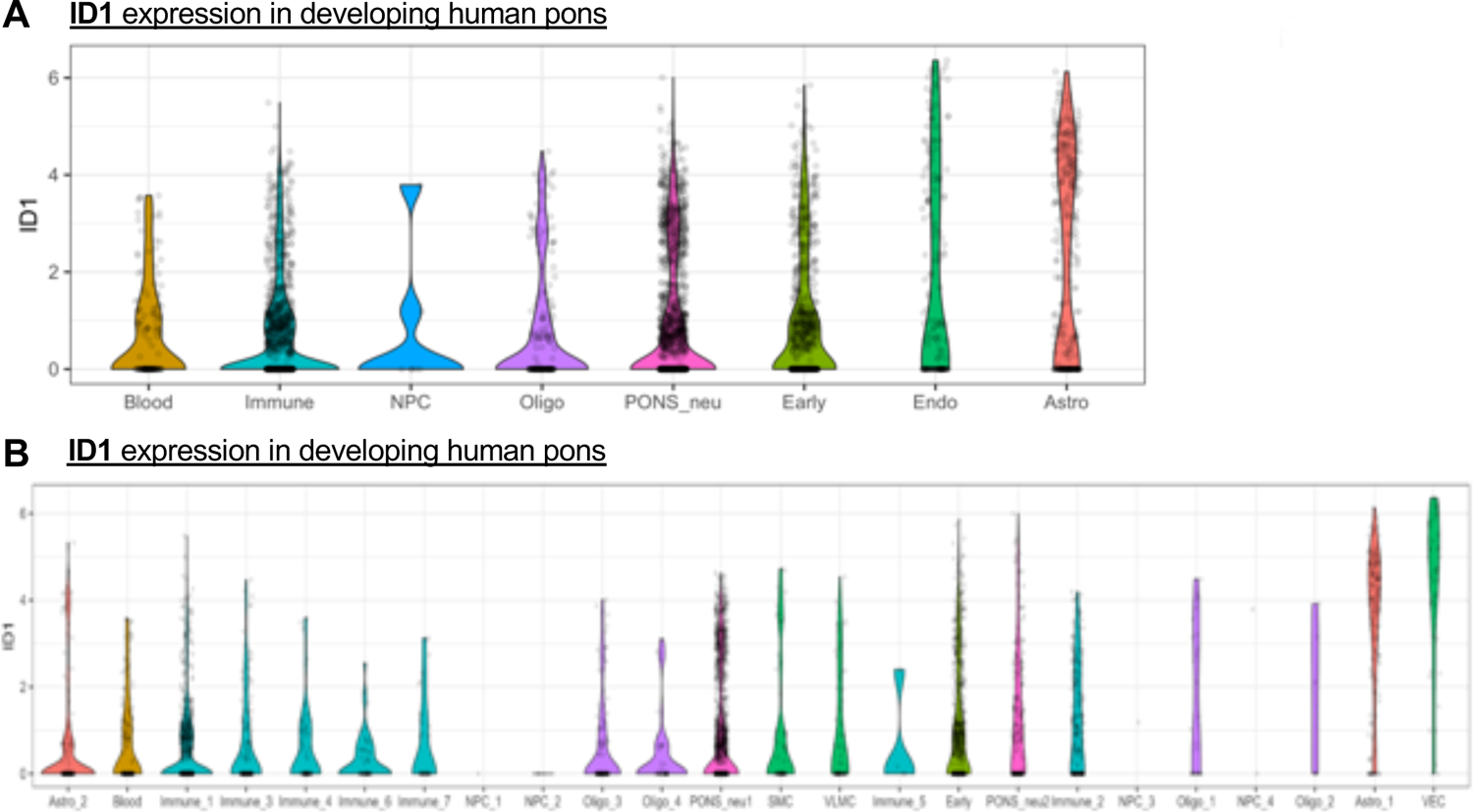
*ID1* expression in varying cell types during normal murine pontine development. **(A-B)** Violin plots from analysis of Fan et al., *Science Advances*, 2020, depicting that AC-like cells show maximum ID1 expression during normal murine pontine development. Data points from all gestational weeks are combined for each cell type and sorted by median. [Astro-Astrocyte].

**Supplementary Figure S7.**
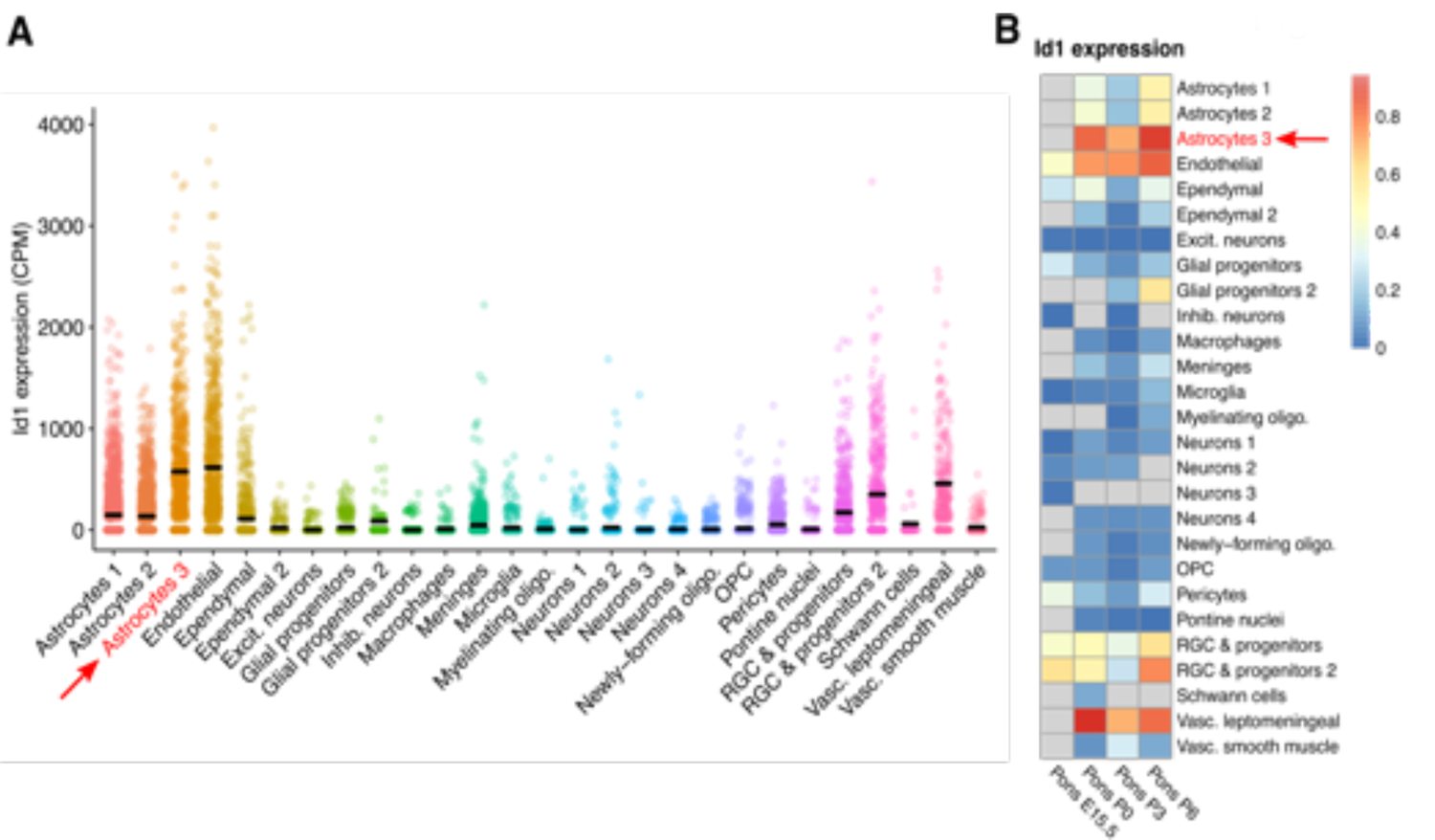
ID1 expression from single-cell transcriptome analysis of varying cell types in normal developing murine pons. **(A)** Single-cell ID1 expression in varying cell types in normal murine pontine development. **(B)** Heatmap of ID1 expression during normal murine pontine development [E15.5-Embryonic day 15.5; P0-Postnatal day 0]. Red arrow indicates increased ID1 expression in astrocytes from P0-P6. Primary data for parts (A) and (B) from Jessa et al., *Nature Genetics*, 2019.

**Supplementary Figure S8.**
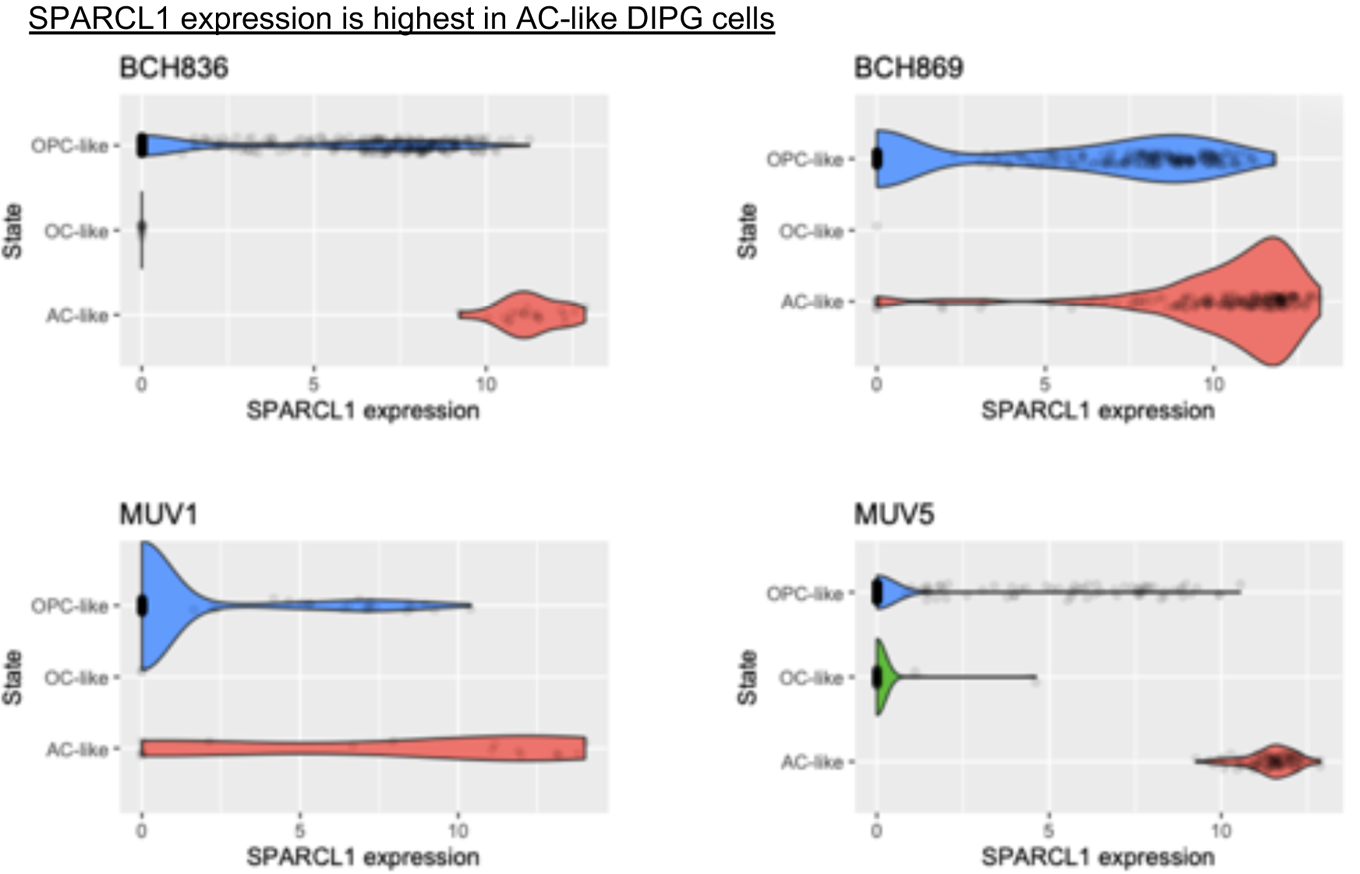
*ID1* expression from single cell RNA-sequencing of four different H3K27M-DMG patients across varying malignant cell types. Violin plots depicting SPARCL1 expression in three subtypes of H3K27M-DIPG malignant cells [Data from pontine DIPG patients in Fig. S1B]. Primary data from Filbin et al., *Science*, 2018. Patients MUV5, BCH836, BCH869-pontine tumors. Patient MUV1-thalamic tumor. [OPC-Oligodendrocyte precursor cell; OC-Oligodendrocyte; AC-Astrocyte].

**Supplementary Figure S9.**
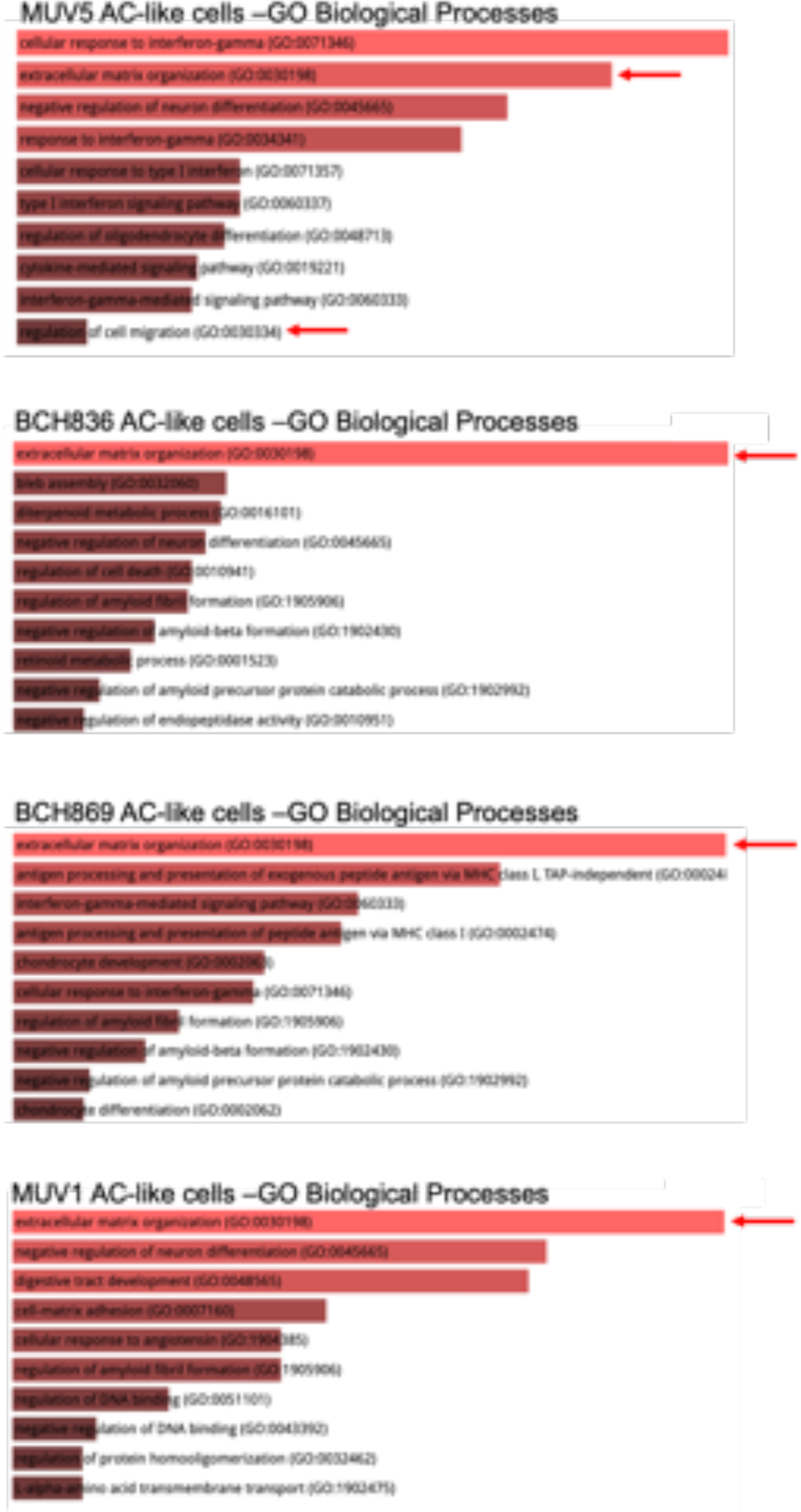
Gene ontology (GO) analysis of higher *ID1*-expressing AC-like cells from H3K27M-mutated tumor patients. GO analysis of primary data from Filbin et al., *Science*, 2018, demonstrates increased expression of genes related to extracellular matrix organization and regulation of cell migration in AC-like cells. Patients MUV5, BCH836, BCH869-pontine tumors. Patient MUV1-thalamic tumor.

**Supplementary Figure S10.**
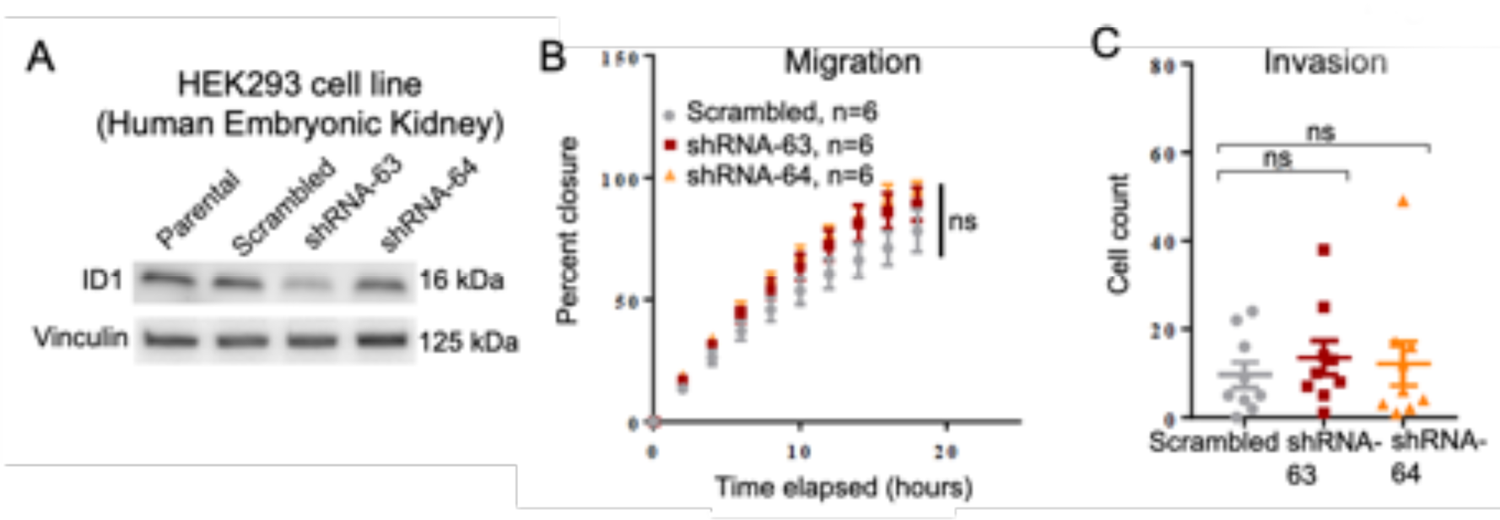
*ID1* knockdown in HEK293 cells. **(A)** Western blot confirming *ID1* knockdown in HEK293 cells. **(B)** Effect of *ID1* knockdown on cell invasion, as measured by Matrigel-coated Boyden chamber assays. Each data point represents an individual image (4 random images were taken per well). NS, P > 0.05, unpaired t test. **(C)** Effect of *ID1* knockdown on migration as measured by scratch assay. NS, P > 0.05, unpaired t test.

**Supplementary Figure S11.**
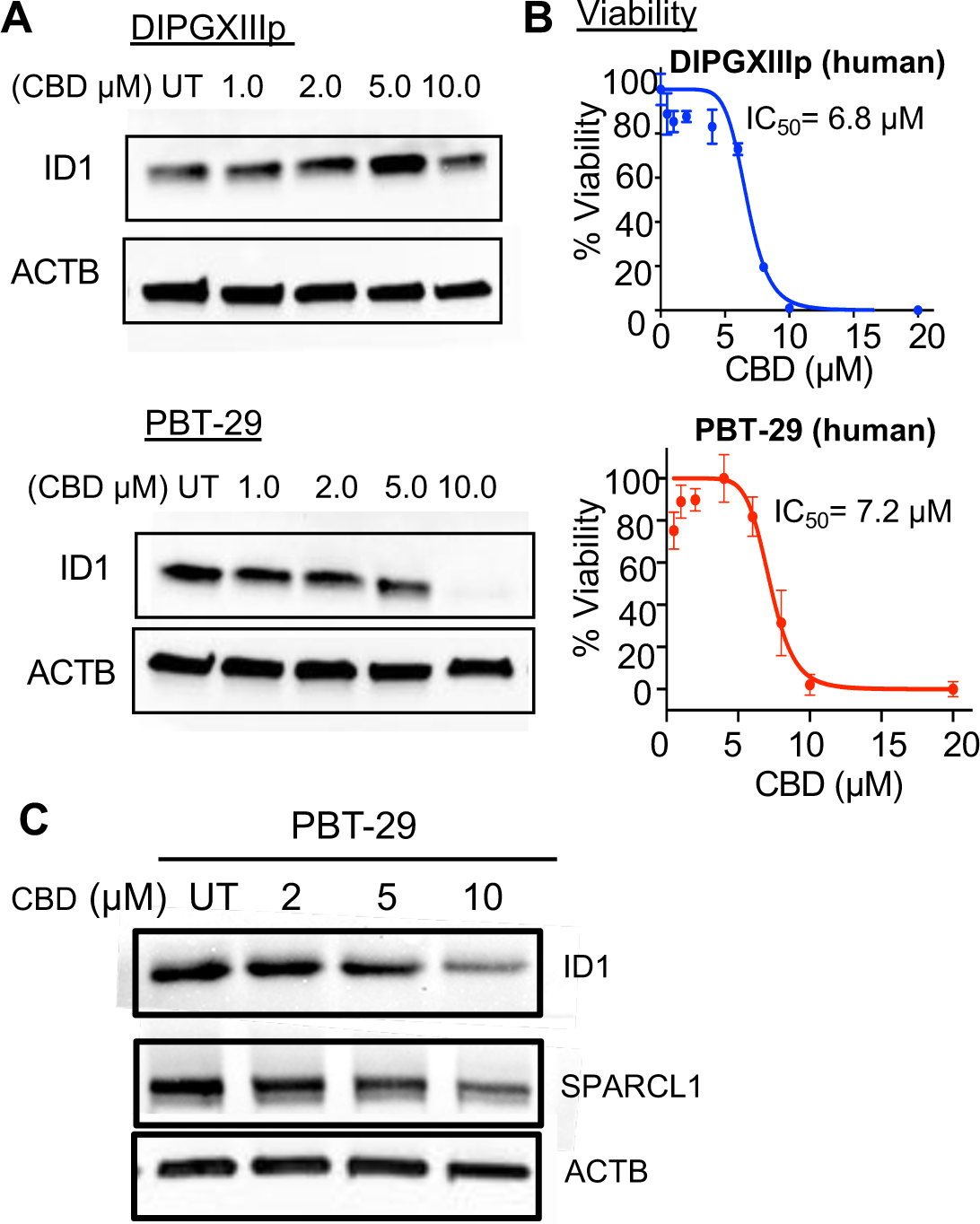
Impact of CBD treatment on ID1 expression in human DIPG cells. **(A)** ID1 western blot of human DIPGXIIp and PBT-29 cells treated with increasing concentrations of CBD, or DMSO control (UT-untreated). Expression levels of ID1 and ACTB were measured. **(B)** Viability of DIPGXIIIp and PBT-029 cells treated with increasing concentrations of CBD (0.5 µM to 20 µM) relative to DMSO-treated control. Experiment was completed in triplicate and data points represent mean +/- SEM. **(C)** Western blot of ID1 and SPARCL1 expression in PBT-29 cells treated with increasing concentrations of CBD or DMSO control (UT). Experiments for all western blots were completed in triplicate.

**Supplementary Figure S12.**
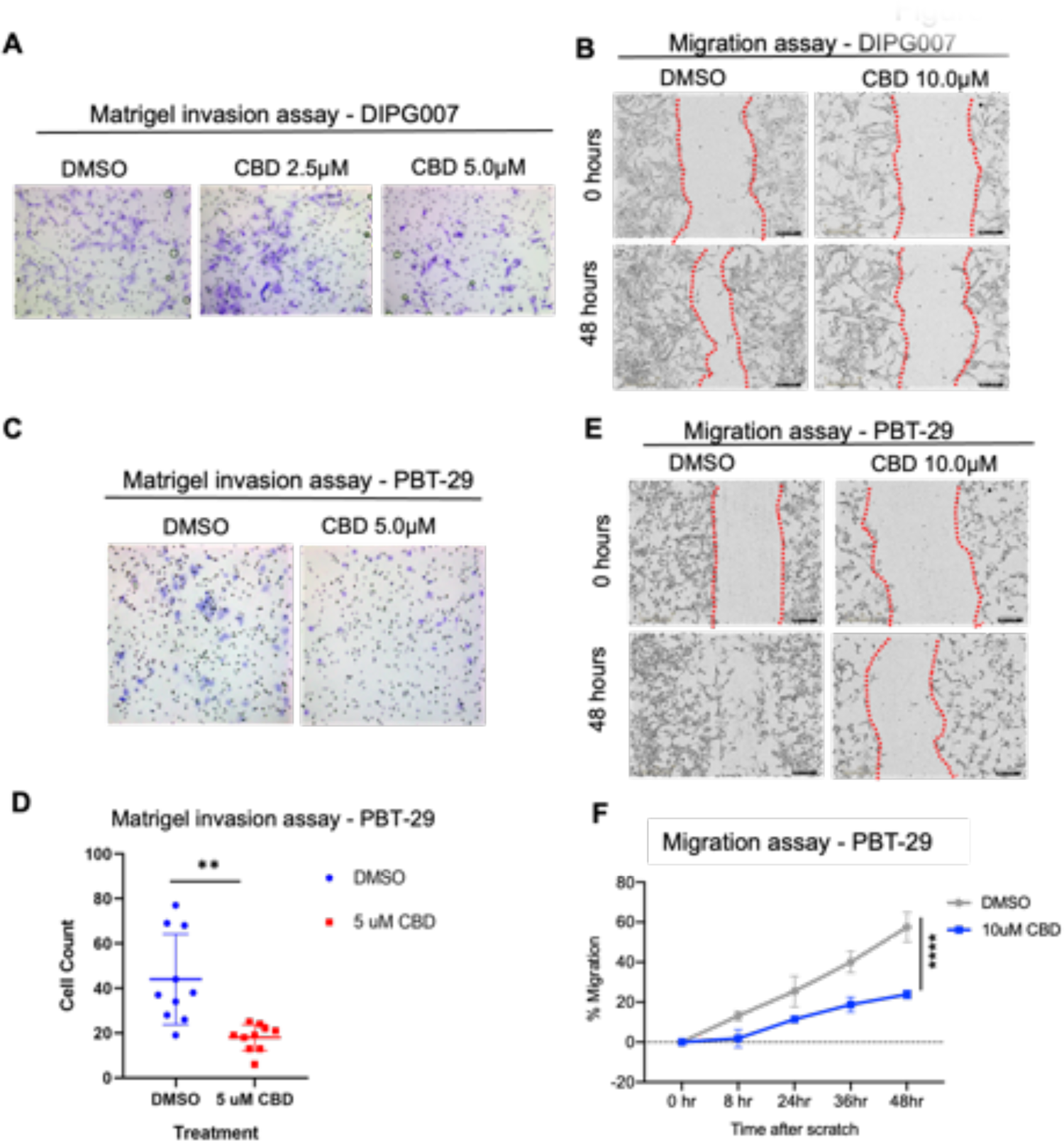
Effect of pharmacologic (CBD) suppression of ID1 on DIPG007 and PBT-29 tumor cell invasion and migration. **(A)** Effect of CBD treatment (2.5µM - 5µM) on invasion of human DIPG007 cells as measured by Matrigel-coated Boyden chamber assay. Images show invading cells stained with crystal violet. Number of invading cells were counted using ImageJ software. **(B)** Images displaying effect of CBD treatment (DMSO control vs 10µM) on DIPG007 cell migration as measured by scratch assay. **(C)** Effect of CBD treatment (5µM) on invasion of human PBT-29 cells as measured by Matrigel-coated Boyden chamber assay. **(D)** Quantification of invading PBT-29 cells treated with either DMSO (control) or 5µM CBD shown in part C determined using ImageJ; **P<0.01, unpaired parametric t test. **(E-F)** Images displaying effect of CBD treatment (DMSO control vs 5µM) on PBT-29 cell migration as measured by scratch assay. Migration was quantified using ImageJ to determine percent wound (outlined with red dashed line) closure. Experiment was completed in triplicate. Data represent mean +/- SEM; ****P<0.0001, unpaired t test. [Magnification for all images is 20x].

**Supplementary Figure S13.**
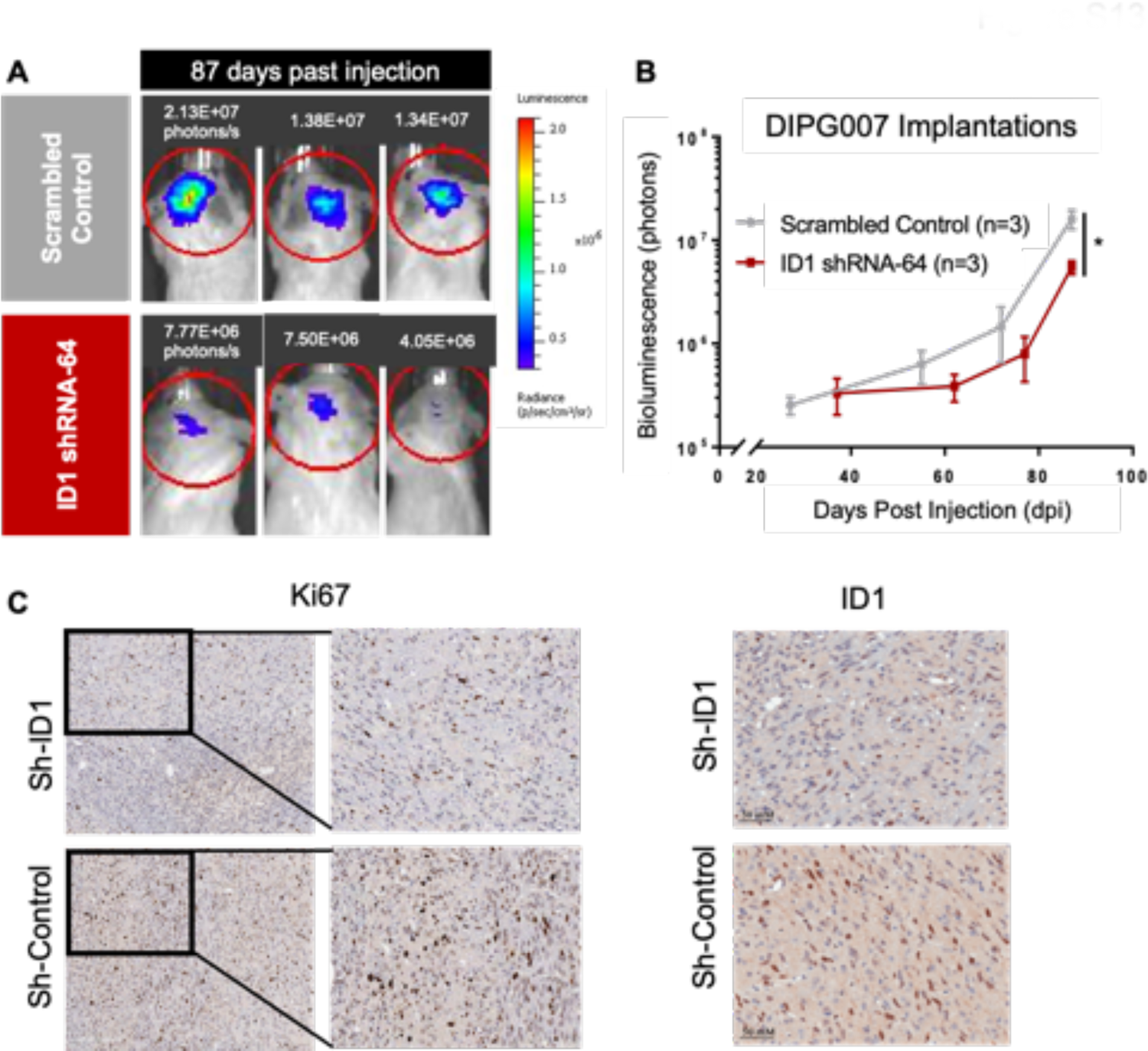
ID1-deficient human DIPG007 cells display slower tumor growth in in vivo model. **(A)** Representative images of bioluminescent tumors from intracranial injection of scrambled-control or ID1-shRNA DIPG007 cells at DPI-97. **(B)** Bioluminescence of intracranially-injected scrambled or ID1-shRNA DIPG007 cells over days-post-injection**. (C)** Example images of IHC staining for Ki67 (left) and ID1 (right) in a sagittal tissue section (tumors generated from implantation of DIPG007 cells). Magnification is 20x.

**Supplementary Figure S14.**
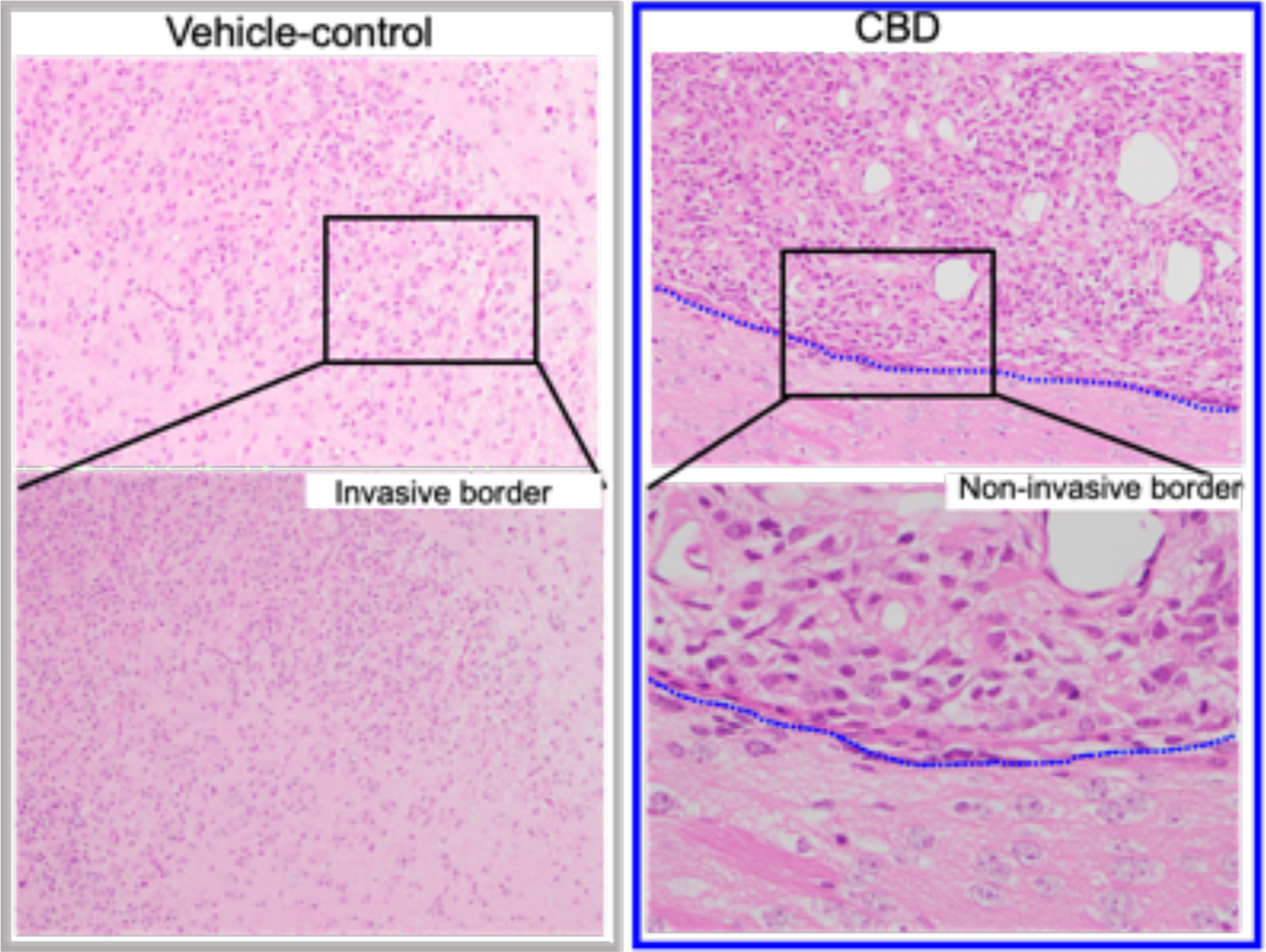
Tumor cell invasion assessment in CBD-vs control-treated PPK mice. Images of IUE-generated PPK tumor borders treated with or without CBD (DMSO vs. 15mg/kg CBD) for assessment of tumor cell invasiveness. Magnification for top row images is 10x and magnification for bottom row is 40x.

**Supplementary Figure S15.**
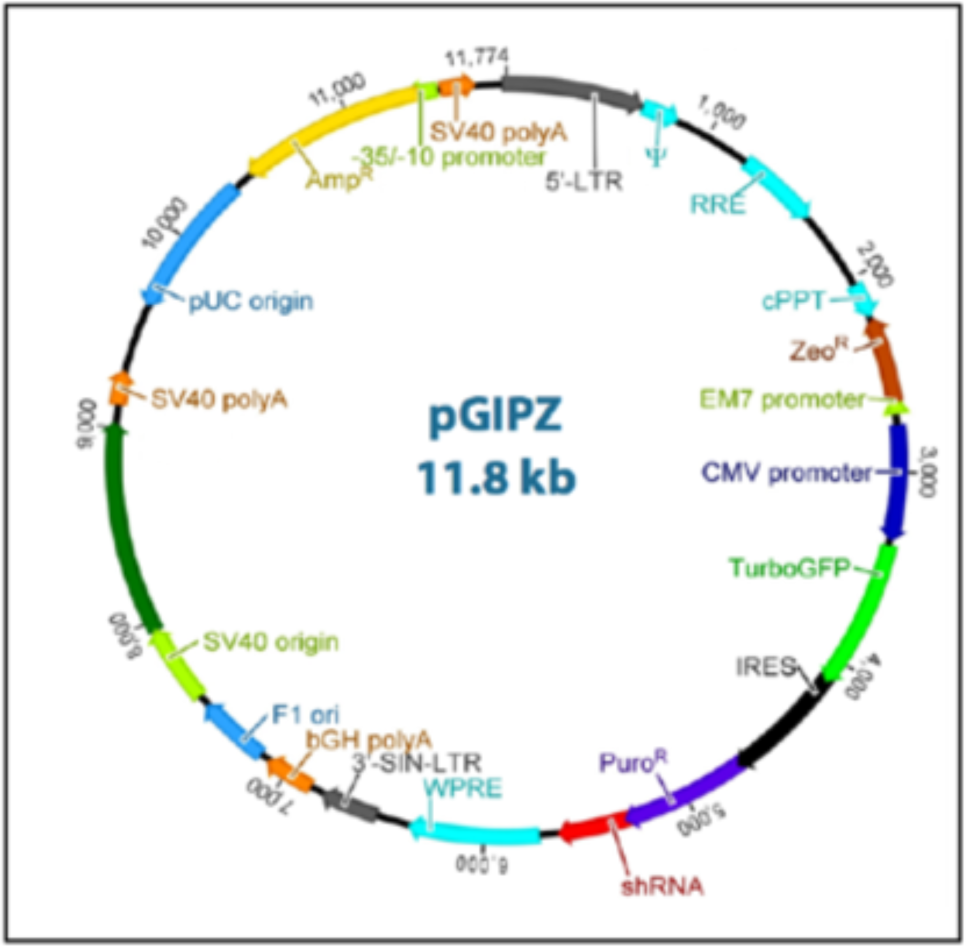
Detailed vector map of pGIPZ lentiviral vector. Lentivirus vector backbone used for ID1-targeting shRNA constructs. Image credit: Dharmacon. Available from: https://dharmacon.horizondiscovery.com/uploadedFiles/Resources/gipz-lentiviral-shrna-manual.pdf

**Supplementary Table S1.**
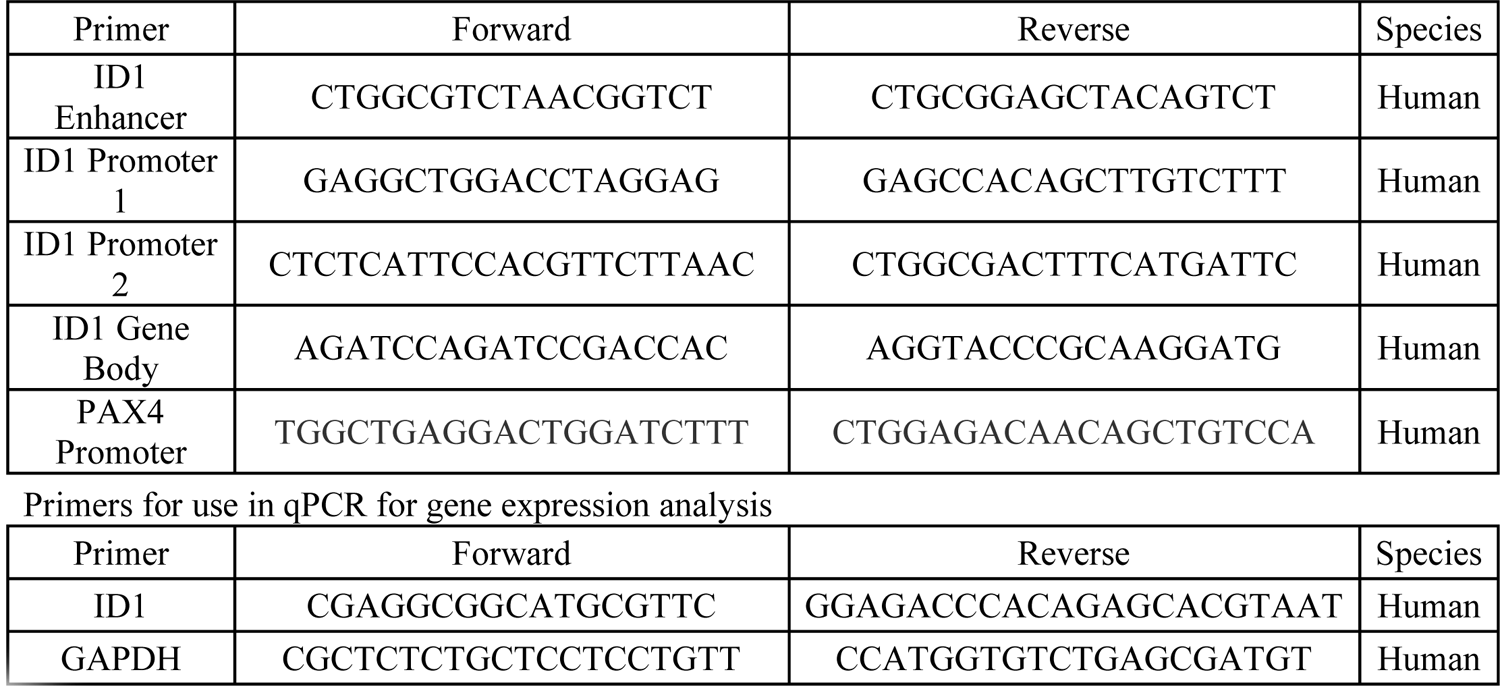
Primer sequences for use in ChIP-qPCR and qPCR rimers for use in qPCR for gene expression analysis

**Supplementary Table S2.** Clinical details of pediatric HGG patients treated with CBD.

| Patient ID | Age | Diagnosis | Molecular | Therapies given | Time to first progression | Overall survival from Diagnosis | CBD duration/dosing/toxicity noted |
| --- | --- | --- | --- | --- | --- | --- | --- |
| CHCO01 | 11 | DMG, H3K27M (thalamus) | H3K27M mutant | Radiation | 24 months | 40 months (still alive) | CBD 2.5g mg (~0.07 mg/kg/day); PO daily; CBD therapy started adjuvantly after initial radiation |
| CHCO02 | 15 | DMG, H3 WT (bi-thalamic) | H3 WT; BARD1 E19* mutation; increased tumor mutational burden | Radiation; TMZ/CCNU; olaparib; pembrolizumab | 14 months | 26 months | CBD 50 mg (0.8 mg/kg/day); PO BID (+THC); CBD therapy started at diagnosis |
| UMPED18 | 9 | DMG, H3K27M (brainstem, DIPG) | H3F3A K27M; PIK3CA E545K; TCF12 V650D | Radiation (+AZD-1775); panobinostat + everolimus | 8 months | 10 months | CBD 300 (13 mg/kg/day) + THC; no toxicity; PO daily; therapy started at radiation – taken until passing |
| UMPED22 | 5 yo | DMG, H3K27M (brainstem, DIPG) | HIST1H3B K27M; ACVR1 G328E | Radiation; hyper-baric O2; Re-irradiation | 9 months | 12 months | CBD 45 mg (2 mg/kg/day) +THC; PO BID or TID; therapy started at radiation – taken until passing |
| UMPED37 | 13 | DMG, H3 WT (bi-thalamic) | H3 WT; EGFR V292L; EGFR in-frame deletion; deletion CDKN2C | Chemotherapy (thioguanine, procarbazine, lomustine, and vincristine); Radiation; osimertinib; bevacizumab | 5 months | 17 months | CBD 150 mg (3 mg/kg/day) after radiation -> 50 mg (1 mg/kg/day), until passing due to nausea; PO daily; therapy started at radiation – taken until passing |
| UMPED56 | 8 yo | DMG, H3K27M (brainstem, DIPG) | HIST1H3B K27M; ACVR1 R206H | Radiation; ONC201; bevacizumab | 13 months | 24 months | CBD (0.6 mg/kg/day) +THC; PO daily; therapy started at radiation – taken until passing |
| UMPED58 | 9 yo | DMG, H3K27M (brainstem, DIPG) | H3F3A K27M; ATRX Q119*; PPM1D G463fs; PDGFRA amplification | Radiation; multi-agent intra-arterial; ONC201 | 18 months | 21 months | CBD (? Dose) + THC; PO or per rectum daily; therapy started at radiation – taken until passing |
| UMPED65 | 16 yo | DMG, H3K27M (brainstem, DIPG) | H3F3A K27M; TP53; PIK3CA | Radiation; ONC201; panobinostat; re-irradiation; paxilisib | 16 months | 24 months | CBD (0.3 mg/kg/day); PO BID/TID, intermittent early in therapy, stopped 7 months prior to passing |
| UMPED67 | 7 yo | DMG, H3K27M (thalamus) | H3F3A K27M; TP53 C277F; NF1 H2434fs | Radiation; ONC201; re-irradiation; bevacizumab | 10 months | 15 months | CBD (0.7 mg/kg/day) + THC; PO daily; therapy started at radiation, stopped 4-5 months prior to passing |
| UMPED69 | 4 yo | DMG, H3K27M (brainstem, DIPG) | HIST1H3B K27M | Radiation; convection-enhanced delivery (CED) trial; re-irradiation; ONC201; paxalisib | 13 months | 28 months | CBD 9 mg (0.5 mg/kg/day) + THC; PO TID; therapy started at radiation – taken until passing |
| UMPED83 | 11 yo | DMG, H3K27M (thalamus) | H3F3A K27M; TP53 S241C | Chemotherapy (temozolomide, irinotecan, bevacizumab); Radiation; ONC201 | 36 months | 60 months | CBD 1500 mg (25 mg/kg/day) + THC; PO TID; therapy started at radiation, stopped one year prior to passing |
| UMPED86 | 7 yo | DMG, H3K27M (brainstem, DIPG) | HIST1H3B K27M; ACVR1 G328E; PI3KCB; PPM1D | Radiation; ONC201; Re-irradiation | 6 months | 8 months | CBD 3 mg (0.4 mg/kg/day); PO TID or QID; therapy started at radiation – taken until passing |
| UMPED97 | 16 | Cortical anaplastic astrocytoma | H3 WT; Tp53 R342* (+germline); CDK4 gain; KRAS gain | Chemotherapy (procarbazine, CCNU, and vincristine); Radiation; irinotecan and bevacizumab | 5 months | 19 months | CBD 400 mg (6 mg/kg/day) after radiation, until passing due to nausea; PO twice daily (Epidiolex); therapy started at radiation – taken until passing |
| UMPED101 | 6 | DMG, H3K27M (brainstem, | H3F3A K27M mutant | Radiation; ONC201; re-irradiation; bevacizumab | 10 months | 14 months | CBD 600 mg (24 mg/kg/day) PO TID; CBD therapy started after radiation – taken until passing |

